# Benchmarking of T-Cell Receptor - Epitope Predictors with ePytope-TCR

**DOI:** 10.1101/2024.11.06.622261

**Authors:** Felix Drost, Anna Chernysheva, Mahmoud Albahah, Katharina Kocher, Kilian Schober, Benjamin Schubert

## Abstract

Understanding the recognition of disease-derived epitopes through T-cell receptors (TCRs) has the potential to serve as a stepping stone for the development of efficient immunotherapies and vaccines. While a plethora of sequence-based prediction methods for TCR-epitope binding exists, their available pre-trained models have not been comparatively evaluated on standardized datasets and evaluation settings. Furthermore, technical problems such as non-standardized input and output formats of these prediction tools hinder interoperability and broad usage in applied research. To alleviate these shortcomings, we introduce ePytope-TCR, an extension of the vaccine design and immuno-prediction framework ePytope. We integrated 18 TCR-epitope prediction methods into this common framework offering interoperable interfaces with standard TCR repertoire data formats. We showcase the applicability of ePytope-TCR by evaluating the performance of the prediction methods on two challenging datasets for annotating single-cell repertoires and predicting TCR cross-reactivity towards mutated epitopes. While novel predictors successfully predicted binding to frequently observed epitopes, all methods failed for less observed epitopes. Further, we detected a strong bias in the prediction scores between different epitope classes. We envision this benchmark to guide researchers in their choice of a predictor for a given setting. Further, we aspire to accelerate the development of novel prediction models by allowing fast benchmarking against existing approaches through common interfaces and defining standardized evaluation settings.

## Introduction

T cells play a fundamental role in the adaptive immune system by recognizing diseased cells through diverse T cell receptors (TCRs). Target cells present antigen-derived peptides – so-called epitopes – bound by the Major Histocompatibility complex (MHC) to the TCR. The TCR mainly interacts with the peptide by the Complementary Determining Region 3 (CDR3) of its *β*-chain while CDR1 and CDR2 ensure contact with the MHC [1]. TCR specificity is crucial for understanding vaccine efficacy [2] and developing immunotherapies against cancer [3] and autoimmune diseases [4]. By better understanding how TCRs recognize specific antigens, we can unlock new strategies for targeted treatments. Therefore, deciphering the TCR-epitope interaction has been declared one of the nine Cancer Grand Challenges in 2023 [5]. Several solutions exist to discover pairs of TCRs and their cognate epitopes such as high-throughput single-cell techniques for staining TCR repertoires with peptide-loaded multimers [6]. However, these experiments are labor- and cost-intensive and typically require an initial set of target epitopes to test. *In silico* prediction methods could overcome these restrictions and provide antigen-specific TCR candidates.

In two seminal papers, Dash et al. [7] and Glanville et al. [7] demonstrated that the TCR sequence is indicative of epitope-specificity when employed for pairwise distance calculation and clustering. Based on these findings, several approaches were developed to compare sequences between a query TCR and an atlas repertoire with known epitope-specificity using string metrics [7, 8], and later deep learning-based representations [9–11]. However, these approaches require the target epitope to be contained in atlases such as the common databases IEDB [12], VDJdb [13], or McPAS-TCR [14], which may not be the case for newly arising epitopes from mutations or novel infectious diseases. With the increasing amount of publicly available TCR-epitope pairs, machine learning models have been trained to predict TCR specificity in two settings. In the first setting, the pre-defined epitopes are considered as categories, i.e., the models receive only the TCR as input and learn from a set of specific TCRs their sequence properties that are characteristic for the epitope class [15, 16]. As they do not incorporate the epitope sequence into their prediction, they can only be applied to the targets on which they were trained. Alternatively, models can encode the epitope sequences as an additional input to learn the TCR-epitope interaction [17, 18]. While such general predictors – in contrast to categorical models – can also be applied to unknown targets, this generalization typically results in a forfeit of predictive performance [19].

While a plethora of prediction models have been published, their comparative performance remains unclear as they have been evaluated on different datasets and under differing settings. Recently, efforts have been made to evaluate methodological development by comparing predictors on standardized datasets [20, 21] notably by the ImmRep workshop [22–24]. However, a thorough benchmark of pre-trained methods is missing to guide immunologists in deciding whether the methods are sufficiently performant for their use case and which of the available models to choose. Additionally, the lack of clearly defined testing standards hinders methodological improvements as the performances of novel methods are difficult to compare. Moreover, the different models are cumbersome to apply from a practical perspective as they employ custom data formats for TCRs and epitopes, discouraging interoperability and limiting their usability for researchers.

We here present ePytope-TCR – an extension to the immune prediction framework ePytope (formerly FRED2 [25]) – to create a simplified interface to TCR-epitope predictors. ePytope-TCR provides a unified framework for 18 pre-trained models allowing their application to TCR repertoires from six common data formats. Additionally, we guide researchers in their tool selection by utilizing ePytope-TCR for a thorough benchmark of these pre-trained models on two challenging datasets focusing on repertoire annotation of single-cell studies [26, 27] and predicting cross-reactivity towards epitope mutations [28].

## Results

### ePytope-TCR provides an extendable interface for TCR-epitope prediction

ePytope (formerly: FRED2 [25]) represents a framework for T cell epitope detection and vaccine design. Its previous implementation (Figure 1a) covered several immunological steps starting from cleavage prediction to MHC-peptide binding prediction, as well as epitope selection and assembly for vaccination. Several external tools can be conveniently combined through interfaces to standardized data representations of transcripts, variants, proteins, peptides, and MHC alleles.

**Figure 1.**
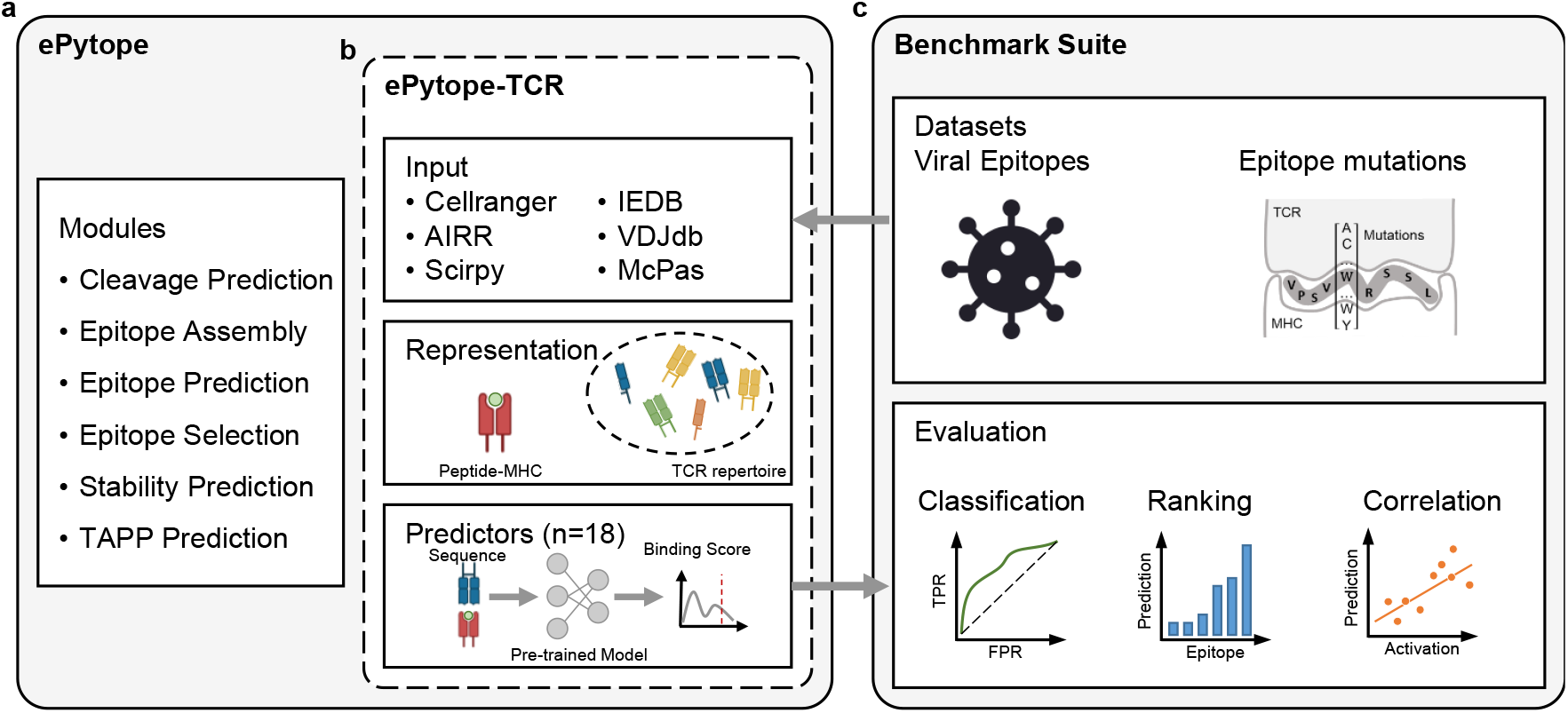
Overview over ePytope-TCR and the benchmark suite. The existing framework of ePytope (**a**) was extended for TCR-epitope prediction (**b**). TCR repertoires can be loaded from six common data formats, and their binding capabilities for pMHCs estimated with 18 sequence-based predictors. **c**, The benchmark suite applies ePytope-TCR to two challenging datasets and evaluates the predictive performance for classification, ranking, and correlation.

To enable the binding prediction between epitopes and TCR repertoires (Figure 1b), we extended ePytope with two data structures. Here, adaptive immune receptors consist of one or several immune receptor chains with known CDR3 amino acid sequences and optional V-, D-, and J-genes in addition to metadata such as T cell type and species. Epitopes represent the combination of peptide sequence and optionally its binding MHC allele through the aforementioned data representations, enabling full interoperability to the remaining framework. To allow users to incorporate their repertoires, ePytope-TCR provides functionality to load TCRs from common single-cell and bulk formats such as the AIRR standard [29], the cellranger-vdj output, and scirpy data object [30]. Additionally, TCRs can be loaded in the formats of the IEDB [12], VDJdb [13], and McPAS-TCR [14] which represent the three most common public databases of TCR-epitope pairs.

Combinations of AIR repertoires and epitopes serve as input for different TCR specificity predictors. While we support 18 predictors for this work (detailed overview in Table 1), novel predictors can be easily integrated by following the defined interface. We utilized the flexibility of ePytope-TCR to evaluate the predictors on two datasets in a standardized fashion via the benchmarking suite (Figure 1c). Overall, ePytope-TCR offers an extendable and easy-to-use tool for applied researchers to predict binding specificity between TCRs and epitope-MHCs. At the same time, it can be used to evaluate new models and make them accessible to the community.

**Table 1.**
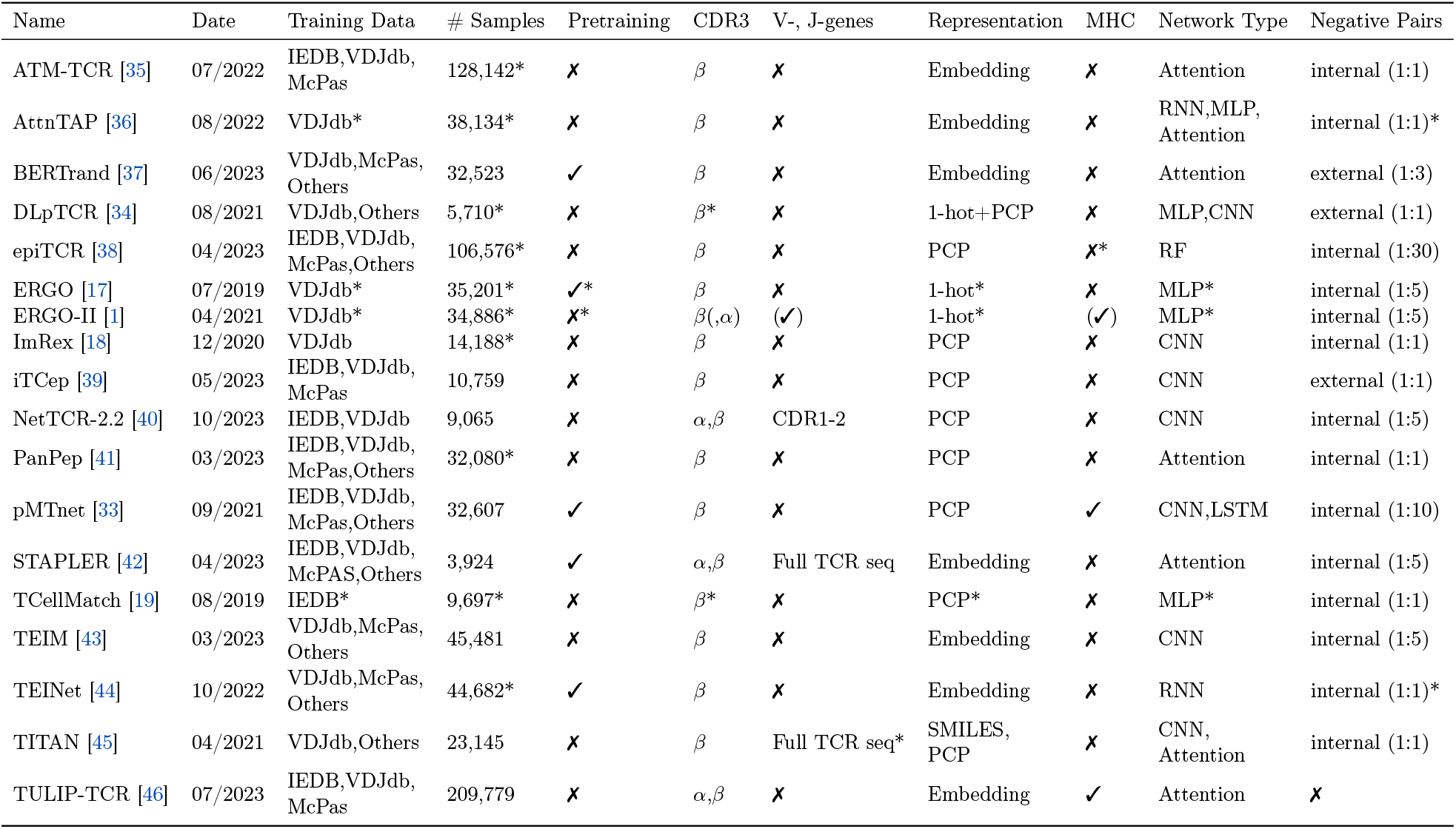
Overview of TCR specificity predictors. The predictors supported by ePytope-TCR are listed with date of publication, model, and training characteristics. The first occurrence in a journal, conference, or preprint is considered the publication date. The table indicates the characteristics of the model version with the highest Area Under the Curve (AUC) in the viral benchmark. If alternative options were additionally available or presented in the publication, the corresponding entry is marked with a star. Optional model inputs are listed in round brackets. # Samples refers to the amount positive training samples. Representation can be either physicochemical properties (PCP), one-hot encoding (1-hot), learned embeddings (Embedding), or SMILES representation (SMILES) [47]. Network type refers to the backbone architecture of the predictor without the classification head consisting of Multi-Layer Perceptron (MLP), Convolutional Neural Networks (CNN), Recurrent Neural Networks (RNNs) such as LSTMs or GRUs, and attention-or transformer-based networks (Attention). Negative data is formed either using the sequences from the matching TCR-pMHC pairs (internal) or by using an additional reference set (external) at a positive-negative ratio of (1:n).

### Predictors

In an extensive literature review (Methods Tool selection), we collected a corpus of 18 sequence-based TCR-epitope specificity predictors (Table 1). As a prerequisite for integration into ePytope-TCR, the pre-trained models must have been publicly available and the models were required to predict arbitrary 9-mer epitopes to allow researchers to apply the method on their own TCR repertoires without retraining the model.

The resulting methods mainly differ in training data, input information, and network style. All models were trained on combinations of the three major databases IEDB [12], VDJdb [13], and McPAS-TCR [14] in addition to smaller datasets [31] or sequencing studies [32, 33]. While all tools required the CDR3*β* amino acid sequence as input, several methods also used the CDR3*α* sequence, the full TCR sequences, CDR1 and CDR2, or MHC type as well. As two exceptional cases, DLpTCR [34] additionally offers a separate CDR3*α*-alone model and ERGO-II [1] takes the *α*-chain, V-and J-gene information, MHC-type, and celltype as optional input. Based on the used databases, the publication date, and the required filtering, the amount of used training data varied drastically between 3,924 and 209,799 positive TCR-epitope pairs. As explained by Dens et al. [21], negative pairs were either created by randomly shuffling the TCR-epitope combinations from the positive pairs (internal), or by matching epitopes to repertoires of TCRs with unknown specificity (external) at ratios from 1:1 to 1:30 between the positive pairs from the databases and these negative combinations.

The majority of models encoded the amino acid sequence either via trained embedding layers or through different physicochemical properties (PCP), while one-hot encodings or other representations were less common. Most of the deep learning models employed a Multilayer Perceptron (MLP) to classify binding versus non-binding, which was often preceded by a feature-extracting network. Here, all common network styles and combinations of them have been tested with attention-based Neural Networks and Convolutional Neural Networks (CNNs) outnumbering Recurrent Neural Networks (RNNs). As two exceptions, epiTCR employed Random Forests (RFs) instead of Deep Neural Networks. Interestingly, TULIP-TCR was not directly trained for classification but in an unsupervised fashion to learn the implicit dependencies between TCR, epitope, and MHC, thereby completely avoiding the need for generating negative training pairs. In a similar direction, several models employed Autoencoding (AE)-or Masked Language Model (MLM)-styled pretraining to learn a TCR representation from a large corpus of TCRs with unknown specificity.

### Strong biases prevail between different viral targets

We investigated to what degree general TCR-epitope predictors can annotate repertoires from sequencing studies. Therefore, we simulated the annotation of a single-cell dataset of 638 TCRs specific to viral epitopes with all 18 predictors through ePytope-TCR. This repertoire was obtained by combining the Severe Acute Respiratory Syndrome Coronavirus 2 (SARS-CoV-2) vaccine study from Kocher et al. [26] with the sample datasets of the BEAM-T pipeline of 10x Genomics [27]. As a ground truth, we utilized the specificity assigned via multimer-staining to one of 14 epitopes from Influenza A virus (IAV), Ebstein-Barr virus (EBV), Cytomegalovirus (CMV), and SARS-CoV-2 bound to five different MHC alleles. As expected in repertoires from sequencing samples, the distribution of TCRs binding to the different epitopes is skewed (Figure 2a) as the four most represented epitopes bind to 76.2% of the TCRs (*n* = 486) while five epitopes are represented less than ten times. We predicted the binding score between each TCR-epitope pair while considering the assigned epitope for each TCR as positive and the 13 other epitopes as negative samples.

**Figure 2.**
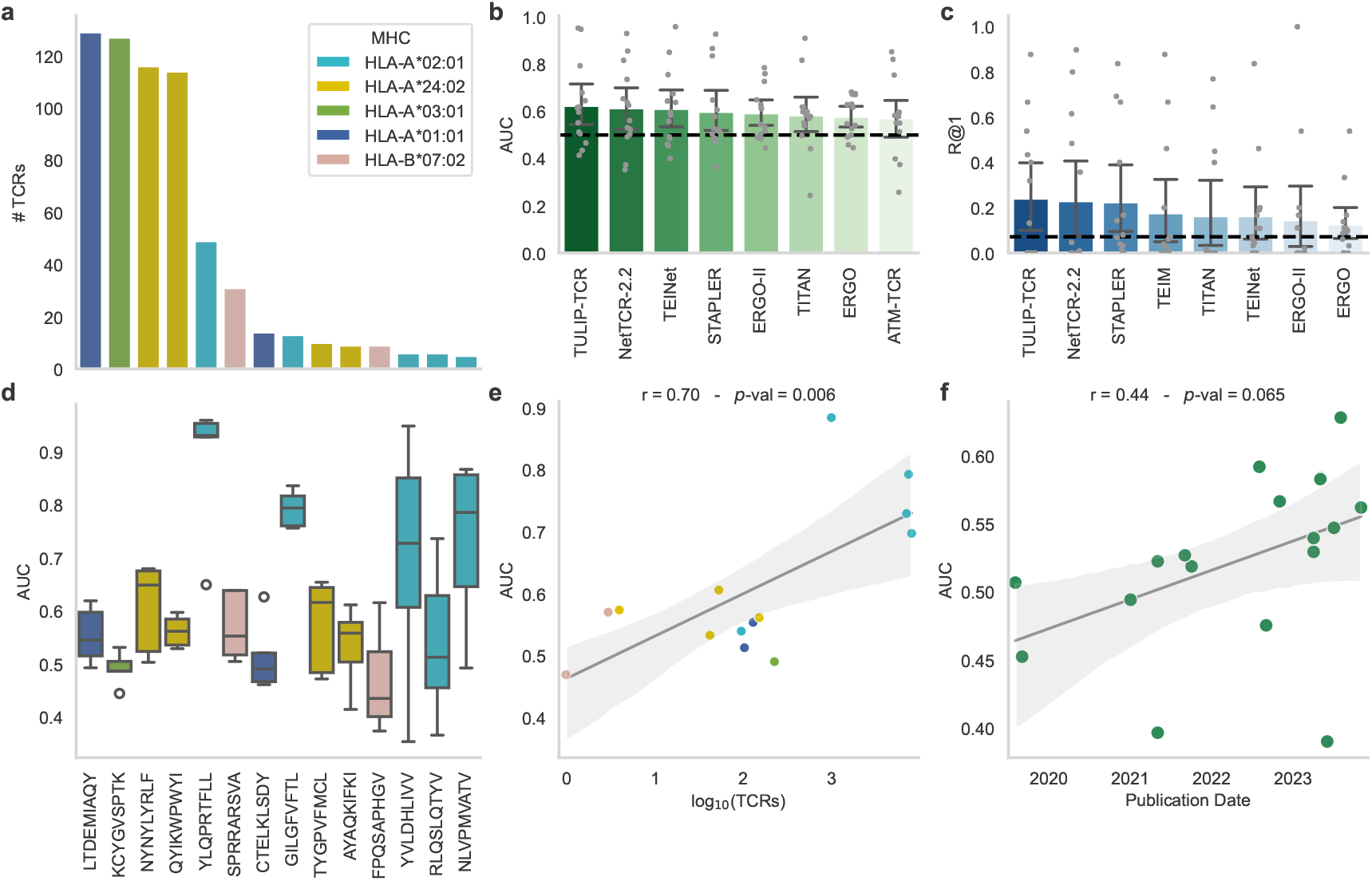
Benchmark on viral epitopes. **a**, The dataset contains 638 TCR-epitope pairs stemming from 14 epitopes, 5 MHC types, and 4 diseases. The best eight predictors measured by the Area Under the Curve (AUC, **b**) and Recall at 1 (R@1, **c**). The mean indicates the average metric score measured for each epitope individually, while the error bars indicate the 95% confidence interval over the scores (*n* = 14 epitopes). The dashed black line marks random predictions. **d**, AUC scores per epitope of the five best performing predictors (*n* = 5 models). The box plot indicates the data quartiles while the median is indicated as a horizontal line. Outliers are marked separately. **e**, Pearson correlation between the average AUC score of the predictors from c and the amount of unique CDR3*β*-epitope pairs available in the combined databases IEDB [12], VDJdb [13], and McPas-TCR [14] over all epitopes (*n* = 14). **f**, Pearson correlation between the average AUC scores and the initial publication date of the method.

In the first step, we evaluated the performance based on Area Under the Curve (AUC) as a common metric for general classification and TCR-epitope prediction. The metric was calculated across the dataset and for each epitope separately (Figure 2b, Supplementary Figure 1a, Supplementary Table 2). Only four methods achieve an average AUC greater than 0.6 with TULIP-TCR performing best at a score of 0.62±0.17 (*n* = 14 epitopes). Eight methods were less than five percent points off the threshold of 0.5 indicating random prediction. Overall, the best-performing predictors remained similar across other classification metrics with a maximal Average Precision Score (APS) of 0.22±0.26 and an F1-Score of 0.16±0.22 (Supplementary Table 2). When a method provided multiple models, we reported the version with the highest AUC in this test and the remainder of this benchmark. Interestingly, the largest differences between model versions were caused by data filtering strategies and choice of dataset for training, with models trained on the VDJdb typically outperforming models trained on McPas-TCR (Supplementary Figure 2a). In practice, the models do not directly classify a TCR-epitope pair as binding or non-binding but provide a continuous prediction score with high values indicating a high binding probability. To assign specificity, practitioners would be required to define thresholds on this prediction score, to decide whether a T cell is specifically binding to an epitope. However, suggestions for binding score thresholds are often not provided in the original publication and are difficult to estimate as the score profile between binders and non-binders overlaps strongly (Supplementary Figure 3a-r). Rank-based metrics offer a more intuitive evaluation in this setting (Supplementary Figure 1b, Supplementary Figure 2, Supplementary Table 2). For each TCR, the prediction scores for the 14 epitopes were ordered, and the position (Average Rank) of the correct match was evaluated with 7.5 as the middle between 1 and 14 indicating random prediction. The Recall at K (R@K) indicates how often the correct epitope occurred within the K highest scores (random for R@1: 1*/*14 = 7.1%). The three methods with the highest score – TULIP-TCR, NetTCR-2.2, and STAPLER – achieved an R@1 between 22-24% and R@3 between 37-38% (Figure 2c). While these predictors exceed the random prediction threshold of 7.1%, applying them for annotation would lead to correct annotation of TCRs in one-fifth to one-fourth of cases, which is insufficient for downstream analysis.

These shortcomings were caused by large discrepancies in performance between the different epitopes, as indicated by a large standard deviation in R@1 greater than 25% and in AUC greater than 15% of the best five predictors (Figure 2b-c, Supplementary Table 2). For five epitopes, at least one predictor reached an AUC greater than 0.75 with a maximum of 0.98 for TEIM on the YLQPRTFLL epitope. Notably, four of these five epitopes were bound by HLA-A*02:01 (Figure 2a), one of the most extensively studied human HLA alleles (Figure 2e). However, for six epitopes, no predictor reached an AUC greater than 0.7. These differences in performance were caused by the over-representation of some epitopes in public databases as indicated by a strong significant Pearson correlation of 0.70 between the average AUC score of the five best predictors and the logarithmized amount of CDR3*β*-sequences in public databases per epitope (Figure 2e). Presumably, much of the performance increase in recent years (Figure 2f) can be attributed to the rise of publicly available TCR-epitope pairs (Figure 4a-b) especially through SARS-CoV-2 epitopes forming the majority of this dataset, while choices in method design had less influence on the performance (Figure 4c). While these performance differences between epitopes clearly hinder the applicability of predictors towards less-observed epitopes, this shortcoming may often be overlooked when calculating metrics across the whole dataset or weighting them by epitope frequency, which will bias the evaluation towards highly-studied epitopes.

Besides large differences in performance, we additionally observed strong biases in the prediction scores between the epitopes. For 13 of 18 methods, we observed a decrease in performance when the AUC was calculated for all epitopes simultaneously compared to the averaged AUC calculated for each epitope individually (Supplementary Figure 5). This indicates that the predictors were better at ordering TCRs within epitopes, but the prediction scores of the models are not comparable between the different epitope targets. This becomes even more apparent when investigating the average prediction score per epitope across the whole dataset (Supplementary Figure 6a). An ideal predictor would assign a score of 1 to all correct binders and 0 otherwise and, thereby, match the epitope frequencies of the dataset. However, 11 predictors have mean binding scores greater than 50% compared to a maximal epitope frequency of 20.2%. This indicates that predictors overestimated binding to certain epitope classes, while presumably less observed epitopes are always predicted as non-binders. This is further displayed by the fact that some predictors show low standard deviation in the prediction scores of all TCRs against an individual epitope, with an average smaller than 10% for four predictors (Supplementary Figure 6b). This indicates that these predictors always provide highly similar prediction scores for a given epitope regardless of the TCR sequence. Presumably, epitopes frequent in the training data were always classified as binding, while others were predicted as non-binding. TULIP-TCR could not be considered in this analysis, as the binding score is not scaled between 0 and 1. However, the next best-performing methods in AUC and R@1 – NetTCR2.2, STAPLER, and TEINet –, all had an overall mean prediction score below 0.2, leading to a profile resembling the true frequencies (Supplementary Figure 6a). Such a comparison between epitopes may be overlooked when evaluating metrics only per epitope. However, prediction scores are required to be comparable across epitopes when annotating repertoires, as otherwise, epitope-specific thresholds need to be defined. Potentially, these shortcomings could be avoided by normalizing against the prediction score of suitable background TCRs which are presumably unspecific to the epitope in question and can thereby be used to estimate the prediction score level.

In summary, our findings suggest that current TCR-epitope prediction tools struggle to accurately assign specificity to viral epitopes in TCR sequencing repertoires in a global manner. While several methods, particularly TULIP-TCR, show strong performance for commonly observed epitopes in public databases, their generalizability to less-represented epitopes is limited, with prediction results nearing random chance. Additionally, significant biases exist between the prediction scores of different epitopes, complicating their comparability between targets. As a result, we recommend using these predictors primarily for epitopes that are well-represented in the common databases, and defining distinct thresholds for each target.

### Epitope mutations are challenging for general specificity predictors

As a second test, we utilized two deep mutational epitope scans from our previous study [28]. This dataset comprised continuous activation scores of six TCRs against 132 single amino-acid mutations of the neo-epitope VPSVWRSSL, and 172 mutations of the human CMV epitope NLVPMVATV (Figure 3a). Predictions on mutational effects could depict potential immune escape of pathogens or identify cross-reactive TCR candidates for cancer immunotherapies. However, prediction of reactivity against epitope mutants represents one of the most challenging tasks for TCR-epitope predictors as the model has to be highly sensitive towards single amino acid changes in the epitope input. Again, we predicted binding with ePytope-TCR for all epitope-TCR combinations and evaluated the AUC based on the activation score threshold defined in the original publication [28] (Supplementary Figure 7a, Supplementary Table 3). TULIP-TCR again performed within the best three predictors at an AUC of 0.59±0.11 (*n* = 26 TCRs) together with TITAN (0.57±0.10) and iTCep (0.61±0.15) (Figure 3b), a method that did not achieve top 8 performance on the viral dataset (Figure 2b-c).

**Figure 3.**
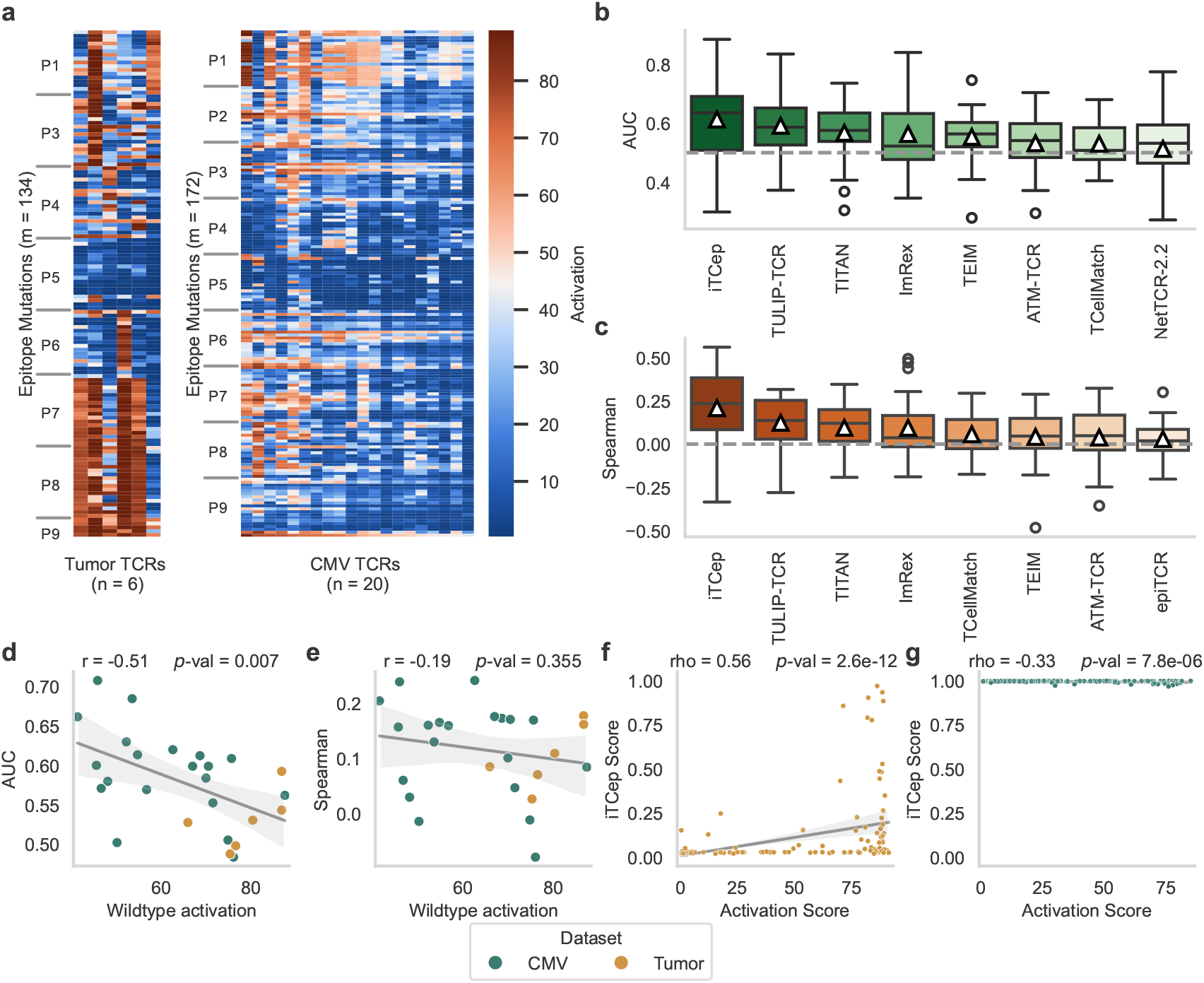
Benchmark on TCR reactivity against epitope mutations. **a**, The dataset consists of two deep mutational scans of TCRs reactive to mutations of the neo-epitope VPSVWRSSL and the human CMV epitope NLVPMVATV. Prediction performance for this dataset is shown for the best eight predictors, measured by the Area Under the Curve (AUC, **b**) and Spearman correlation coefficient (Spearman, **c**). The box plot indicates the data quartiles while the median is indicated as a horizontal line and the mean as a white triangle (*n* = 14 TCRs). Outliers are marked separately. The dashed black line marks random predictions. Pearson correlation (*n* = 26 TCRs) between the average AUC (**d**) or Spearman coefficient (**e**) and the activation score to the wildtype epitope. Each point represents one TCR with the performance scores averaged for the best predictors (*n* = 5) on the corresponding metric from b and c, respectively. Spearman correlation between the prediction score of iTCep and the activation score for all epitope mutations for two selected TCRs with the highest (**f**, *n* = 133 epitopes) and the lowest (**g**, *n* = 172 epitopes) correlation score.

The data on mutational epitope scans additionally provide a continuous score for each TCR-epitope pair relating to the fraction of T cells from this clonotype that were activated by a mutated epitope peptide. As a next step, we, therefore, evaluated to what degree the models’ prediction scores correlate to the respective activation scores in order to investigate to what degree continuous T cell activation was captured by the predictors. The overall performance largely followed the classification setting (indicated by AUC in Figure 3b) on this dataset and was with a maximum of 0.21±0.22 in Spearman rank coefficient close to the random prediction boundary of 0.0 (Figure 3c, Supplementary Figure 7b, Supplementary Table 3). Interestingly, a variant of TCellmatch – one of the first general epitope-TCR predictors – achieved the highest AUC (0.62±0.11) and Spearman coefficient (0.29±0.16) on this dataset (Supplementary Figure 8) but is not listed in the results as it is not the version selected in the previous evaluation (Supplementary Figure 2). These findings suggested that none of the methods so far is suitable for predicting the effect of point mutations in epitopes.

As in the viral dataset, we observed large differences in performance between the different TCRs and targets. We observed a significant negative Pearson correlation (r=-0.51) between the average performance scores of the best five predictors in AUC (Figure 3b) to the activation score of the TCRs (*n* = 26) against the wildtype epitope (Figure 3d). The correlation (*ρ* =-0.19) of the best five models (Figure 3c) was not significant for the Spearman coefficient (Figure 3e). Further, the average performance scores of the five best predictors decreased for neo-epitope reactive TCRs compared to CMV reactive TCRs at an AUC of 0.06. Again, we observed a strong bias between the different target epitope variants with regard to the continuous prediction scores that the models provide as an output to indicate binding. Even for the two leading methods, iTCep and TULIP-TCR, the prediction scores between both base epitopes separated into a clear bimodal distribution (Supplementary Figure 9, Supplementary Figure 10). 14 of the 18 tested methods yielded lower average prediction scores for the tumor epitope mutations despite them surpassing the activation threshold by 38.9% more often than the CMV epitopes. As NLVPMVATV is one of the epitopes with the most recorded TCRs in public databases, prediction methods were inherently biased to assume binding with a higher probability compared to the unobserved VPSVWRSSL neo-epitope which is reflected by a higher level of prediction scores. This can be illustrated well in the example of the best-performing method iTCep. The TCR with the highest Spearman coefficient of 0.56 (R25, tumor) showed variable prediction scores between 0 and 1 (Figure 3f). While many mutations would be incorrectly classified as non-binding based on low prediction scores, several epitopes show higher scores indicating that the model successfully incorporated the epitope sequence into its prediction. In contrast, all mutations are predicted to bind for the TCR with the lowest coefficient of -0.33 (CMV81-14, CMV) at a prediction score greater than 0.95 with a standard deviation of 0.005 (Figure 3f). Apparently, the model correctly predicted the binding of the TCR towards the base epitope but failed to take the negative effects of mutations into account, resulting in high prediction scores for all epitopes. This again shows the biases different epitope targets imprint on the prediction score based on their prevalence in public databases.

In summary, general TCR-epitope predictors cannot reliably predict the effect of epitope mutations. This highlights the need for specialized datasets and prediction models such as P-TEAM, which was proposed by us recently [28]. We observe strong biases between different target classes highlighting the data dependencies of current predictors. Of the best models in the viral dataset, TULIP-TCR is also ranked high in the mutation dataset. While no current model can be utilized for practical use in this setting, we hope that this benchmark spawns research interest and we expect a large potential for predicting epitope reactivities in future works.

## Discussion

*In-silico* binding prediction of a TCR towards pathogen-, tumor-, or self-derived epitopes will signify a pivotal step in computer-aided vaccine and immunotherapy development as well as for the investigation of TCR repertoire evolution. While several methods have been proposed to predict specificity based on the TCR and epitope sequences, they have not yet found wide application for immunological research. On the one hand, these proposed approaches widely lack interoperability, among themselves and to standard repertoire data formats. On the other hand, their predictive performance in applied use cases has remained unknown. Despite notable efforts by the ImmRep Workshop [22–24], many pre-trained models of the predictors have not been systematically evaluated under benchmark conditions, and no defined testing rules and datasets exist to guide the development of new methods.

To alleviate these shortcomings, we here introduce ePytope-TCR as a TCR-epitope prediction extension of the vaccine design and immuno-prediction framework ePytope (formerly FRED2 [25]). We incorporated a unified interface to 18 pre-trained TCR-specificity predictors and import utilities to three common databases and three standardized repertoire data formats. We showcase the capabilities of ePytope-TCR by applying the predictors to two benchmarking datasets, which test their ability to annotate single-cell datasets with specificity towards viral epitopes and to predict the cross-reactivity of TCRs against epitope mutations.

Despite overall low metric scores for classification and ranking when averaged across targets, we found sufficient performance for epitopes with large support in public databases with AUC scores greater than 75% on five of the tested epitopes. These results are in agreement with previous findings, that TCR-epitope prediction is still limited by the lack of diverse training data and predictors fail for unseen targets [20, 48]. However, we observed increasing performance in recent models which may partially attributed to the rise in available TCR-epitope pairs. Therefore, we advise applying these predictors only for target epitopes covered in the methods’ respective training data. In particular, TULIP-TCR scored within the best three models measured by the AUC and R@1 in the viral dataset and AUC and Spearman in the mutation dataset.

While the predictors were capable of annotating specificity for selected epitopes, they were not able to reliably predict the effect of mutations in known or unknown wild-type epitopes on T-cell function. We hope this benchmark encourages further interest in this use case as one of the most challenging tasks in TCR-epitope prediction.

In both datasets, we observed a strong bias in the prediction scores dependent on the target epitope. This further complicates applying the predictors for annotation as they therefore require epitope-specific classification thresholds. Both, the different performance between epitopes and the bias in prediction scores, highlight important aspects for the evaluation of TCR-epitope predictors. On the one hand, the metric scores are stirred towards frequently observed epitopes and low performance on rare epitopes might not be discovered when metrics are evaluated across datasets or weighted by class support. On the other hand, biases in prediction score levels might not be discovered when evaluating only within epitope classes. Therefore, we advise performing both evaluations separately to allow a holistic view of the models’ performance.

This evaluation provides an overview of the performance of current general TCR-epitope predictors. However, our tests are – similar to the methods themselves – limited by the data at hand. First, we only evaluated predictions for CD8^+^/MHC-I epitopes as this is the major focus of current research. To enable a complete evaluation between all predictors, we disregard all non-9-mer epitopes leading to a limited amount of 638 TCRs in the viral dataset and two scans in the mutation dataset. Further, the epitopes were bound by only five different MHC classes which did not allow us to observe strong biases between alleles independent from the amount of reported TCRs. Additionally, all epitopes of the viral dataset were observed in public databases at least once. An improved benchmark would ideally contain several epitopes without any reported binding TCRs to evaluate generalization for out-of-target predictions. Despite our best efforts, we also cannot rule out a small amount of wrongly annotated TCRs in the viral benchmark as multimer-staining and the resulting annotation may be susceptible to background noise. Finally, in this work, we evaluated the performance of pre-trained models and did not retrain any model ourselves. Therefore, we cannot separate whether differences in performance were caused by the methods’ design choices or by their underlying training data. A standardized test of different architectures, training paradigms, data filtering, TCR information, and pre-training strategies remains open for future research.

Our contribution to TCR-epitope prediction in this work is two-fold. First, we make TCR-epitope prediction methods available for the applied community by providing an interoperable interface with ePytope-TCR, which allows researchers to apply the predictors to their own repertoire data. Through the benchmark, we further guide them in which settings the models can be applied, which mainly reduces them in settings where the epitopes have been well-researched. Second, we provide the basis for accelerating future method development by defining evaluation datasets and metrics and allowing fast benchmarking to other methods through ePytope-TCR. Overall, we envision improved evaluation and interoperability will finally bridge the gap in applying TCR-epitope predictors to relevant immunological studies of large-scale TCR repertoires, vaccine development, and identifying therapeutic TCR candidates against pathogens and tumors.

## Code availability

All source code to reproduce the results of this manuscript are publicly available. The software package ePytope-TCR including tutorials is available at https://github.com/SchubertLab/epytope. The benchmarking suite including the source code to reproduce the results and figures of the evaluation can be accessed at https://github.com/SchubertLab/benchmark_TCRprediction. The repositories of the individual predictors were linked in Supplementary Table 1.

## Data availability

All datasets used in this manuscript are publicly available and provided through the benchmarking suite. The unprocessed data of the SARS-CoV-2 can be accessed via Zenodo. The raw sequencing data of four samples from 10x Genomics can be obtained on the company’s homepage (S1, S2, S3, S4). The unprocessed data from the mutation dataset can be accessed in the Supplementary Material of the origin publication [28].

## Author Contributions

F.D. and B.S. conceived the project and supervised the research. F.D., A.C., and M.A. integrated the models. F.D. and A.C. performed the literature review. F.D. supervised the integration and implemented and performed the evaluation. K.K. and K.S. generated the viral test data and provided it prior to publication. K.S. provided feedback on the evaluation and the manuscript. All authors wrote the manuscript.

## Acknowledgments

This work was supported by the BMBF grant DeepTCR (#031L0290A) and the de.NBI Cloud within the German Network for Bioinformatics Infrastructure (de.NBI) and ELIXIR-DE (Forschungszentrum Jülich and W-de.NBI-001, W-de.NBI-004, W-de.NBI-008, W-de.NBI-010, W-de.NBI-013, W-de.NBI-014, W-de.NBI-016, W-de.NBI-022). F.D. is supported by the Helmholtz Association under the joint research school “Munich School for Data Science -MUDS” and acknowledges financial support from the Joachim Herz Stiftung.

## Material and Methods

### Tool selection

We conducted a literature review by examining ERGO [17], NetTCR [16], TCellmatch [19], and TITAN [45] as examples of the first generation of general TCR-epitope predictors. Next, we reviewed the references of each publication to identify prior work on this topic. Based on this initial list, we included all prediction-related publications that referenced any of these methods through an iterative process. After composing this comprehensive list in November 2023, we excluded models that did not meet the requirements of this benchmark. Specifically, we removed predictors that treated the epitope as a category or performed atlas-query mapping, as these approaches are not suited for general TCR-epitope prediction. All models were required to make predictions at the sequence level without relying on structural information or modeling. Additionally, the source code and pre-trained model weights had to be released under a license open for academic use.

### Predictors

If not stated otherwise, the predictors were integrated into ePytope-TCR based on their original implementation as described in their code repository (Supplementary Table 1). However, several predictors required minor adjustments listed in the following for reproducibility.

#### pMTnet

In contrast to the other models, pMTnet is trained to indicate a high binding probability by a low score instead of a high score. We, therefore, subtracted the resulting score from one to keep the score direction uniform within ePytope-TCR with high scores indicating a high binding probability.

#### ERGO-II

To execute the predictor, the original implementation was slightly modified by ePytope-TCR. The model was changed to automatically select the available execution device, and the path to the pre-trained models was changed to point to the correct directory.

#### TULIP-TCR

Similar to ERGO-II, the execution device was adjusted and unspecified program arguments were exchanged. Additionally, the quantitative score and the used MHC were appended to the output of the predictor.

#### STAPLER

STAPLER requires full TCR sequence information. ePytope-TCR, however, provides the CDR3 sequence in addition to the categorical label of V-, D-, and J-gene, as this is the format provided by most databases. Therefore, Stitchr [49] was used to recreate the full TCR sequence.

#### NetTCR2.2

The full TCR sequence was obtained as described for STAPLER. Following, the CDR1 and CDR2 sequences for *α*- and *β*-chain were identified with ANARCI [50] as described in the publication [40].

#### DLpTCR

The *α*-*β* version of DLpTCR was excluded from the benchmark as it does not provide a continuous prediction score, but rather a binary label, which is not suitable for the performance metrics chosen in this benchmark.

### Datasets

The two datasets utilized in this benchmark were publicly available, but not yet incorporated into public databases which was important to avoid data leakage to the models’ training data. The initial data were preprocessed by the benchmarking suite to facilitate a standardized evaluation. Generally, we eliminate TCRs with unknown specificity and missing annotation of CDR3 sequences, V-, or J-genes on *α*-and *β*-chain. To maintain consistency in the evaluation set, TCRs with CDR3 length greater than 19 were excluded, along with non-9-mer epitopes, as some predictors are restricted to 9-mer epitopes.

#### Viral datasets

We combined two single-cell datasets stemming from a Severe Acute Respiratory Syndrome Coronavirus 2 (SARS-CoV-2) vaccine cohort study by Kocher et al. [26] and the BEAM-T pipeline example datasets provided by 10x Genomics on their company’s website. Both datasets include TCR sequencing and staining with DNA-barcoded peptide-MHC multimers to assign epitope specificity. The vaccine cohort contains TCRs reactive to SARS-CoV-2 epitopes as well as other viral controls to Human Herpes Virus 1 (HHV-1), Influenza A Virus (IAV), and Epstein-Barr Virus (EBV) from 14 donors including a hypervaccinated individual, who at that point has received 215 separate vaccinations [51]. Peripheral blood mononuclear cells (PBMCs) were sequenced for transcriptome, TCRs, and partially surface protein markers. The specificity was identified through sample- and epitope-specific thresholds on UMI counts of peptide-loaded MHC-dextramers. The underlying annotated data object can be obtained from Zenodo, from which we collected clonotypes to all 9-mer epitopes described in the publication. For the BEAM-T data [27], the raw sequencing data for four samples were downloaded from the 10x Genomics website (accessed April 2024) which contain single-cell transcriptome, TCR, and multimer-staining against eleven viral epitopes from human Cytomegalovirus (CMV), EBV, IAV, and SARS-CoV-2. The raw data were processed using the ‘cellranger multi’ command with cellranger (7.1.0). All cells without identified TCRs were removed, and specificity was assigned based on the BEAM algorithm’s score at a threshold of 92.5% as suggested by 10x Genomics. TCRs that were cross-reactive based on this definition either on a cell-or on a clonotype-level were removed from the dataset. Non-9-mer epitopes and epitopes bound by less than five TCRs in the joint dataset were removed in addition to 48 TCR-epitope pairs which were already contained in IEDB [12], VDJdb [13], or McPAS-TCR [14]. Finally, the combined viral dataset resulted in a total of 638 TCRs which were reactive to one of 14 epitopes bound to one of five different MHC alleles.

#### Mutation dataset

This dataset originated from our previous publication [28], in which we studied and predicted the effect of epitope mutations on T cell activation. The dataset comprises three TCR repertoires with annotated activation scores toward deep mutational epitope scans, i.e., each epitope residue was systematically exchanged by all 19 other amino acids. Details on data acquisition can be found in the original publications [28, 52]. In short, the TCRs of each repertoire were expressed in Jurkat triple parameter reporter cells (JTPRs) and co-incubated with peptide-pulsed splenocytes. For each combination of mutated epitopes and TCRs, a continuous activation score was assessed through NFAT reporter expression in flow cytometry. As the many predictors were predominantly trained on human TCRs and MHCs, we selected the two human datasets, which contain TCRs against the neo-epitope VPSVWRSSL and the human CMV epitope NLVPMVATV. Following the original publication, the activation score was binarized by the thresholds of 66.09% and 40.0% for the neo-antigen and CMV datasets, respectively, representing active T-cell recruitment and activation. TCR-pMHC pairs of the TCR R27 were excluded from the neo-antigen dataset as no peptide surpassed the binarization threshold. The corresponding HLA alleles *HLA-B*07:02* for the neo-epitope and *HLA-A*02:01* for the CMV epitope were added as an annotation to each TCR-epitope pair. As all peptides stem from single point mutations of the 9-mer epitopes and the TCR CDR3 sequences lie within the allowed ranges of all predictors, no additional TCR-pMHC pairs were filtered.

Over time, these datasets may be added to public databases of TCR-epitope pairs. For the viral dataset, we made sure that no CDR3-epitope pair was present in the database during the development of the predictors. Although some TCR-wildtype epitope pairs from the mutation dataset may have been part of a training dataset for the selected prediction model, none of the CDR3 and mutated epitope pairs are included in any of the commonly used TCR-specificity databases. We propose that other methods evaluated with the benchmarking suite carefully filter their training and validation sets for TCR-epitope pairs following the rules outlined here.

#### Metrics

All predictors provide a continuous score to indicate the binding of a TCR to an epitope, where higher scores imply a higher chance of binding. To evaluate the models, we employed classification metrics (AUC, APS, F1-Score), rank-based metrics (Recall at K, Average Rank), and correlation-based metrics (Pearson correlation, Spearman rank coefficient). For the classification metrics, a binding TCR-epitope pair was considered as a positive sample. For the viral dataset, all other possible combinations of TCR-epitope were considered negative samples. All samples not exceeding the binarization thresholds in the mutation dataset were considered non-binding. If not indicated otherwise, we refer to the metrics calculated within groups, i.e., per epitopes in the viral dataset and per TCR in the mutation dataset. Additionally, we report the metrics macro-averaged across the whole dataset, i.e., each class in the group has equal weight.

##### AUC, APS

The Area Under the Receiver Operating Characteristic (AUC) and Average Precision Score (APS) both summarize the prediction quality across all classification thresholds indicating how well the predicted value separates positive from negative pairs. While the AUC calculates the area beneath the curve of Recall vs False positive rate, the APS utilizes the Precision-Recall curve.

##### F1-Score

The F1-Score represents the harmonic mean between recall and precision. To determine the classification boundary, the F1-score on all prediction values was evaluated individually for each predictor and dataset, and the threshold resulting in the highest value was chosen.

##### Recall at K (R@K)

Given a repertoire of TCRs that are known to bind to a limited set of epitopes, rank-based metrics describe how well the correct epitope can be identified for each TCR. Following its definition as described in [53], we consider the R@K as the average of how often the correct epitope is contained in the top K predictions for an individual TCR.

##### Average Rank

In the same setting as for R@K, the set of epitopes is ordered by prediction value. The Average Rank corresponds to the mean position of the cognate epitope within this order over all TCRs.

##### Pearson and Spearman correlation

While the other metrics evaluate the predictors’ classification capability, correlation-based metrics indicate whether the prediction score is also quantitatively associated with TCR binding or activation. While the Pearson coefficient measures the linear relationship between prediction and the true score, the Spearman rank coefficient describes their potentially non-linear monotonic relationship, i.e., comparing the orders within both scores.

## Supplementary Figures

**Supplementary Figure 1.**
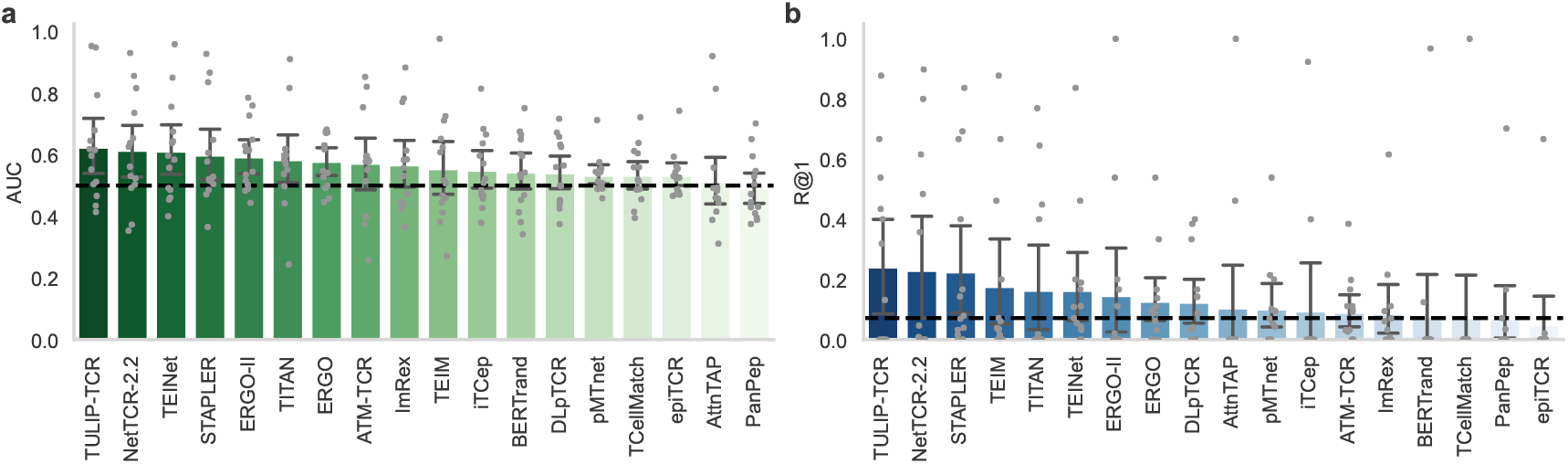
Performance of all models on the viral dataset. Performance scores of all models on the viral benchmark for AUC (**a**) and R@1 (**b**). The models are ordered by their performance on the respective metric. The mean indicates the average metric score measured for each epitope individually, while the error bars indicate the 95% confidence interval over the scores (*n* = 14 epitopes). The dashed black line marks random predictions.

**Supplementary Figure 2.**
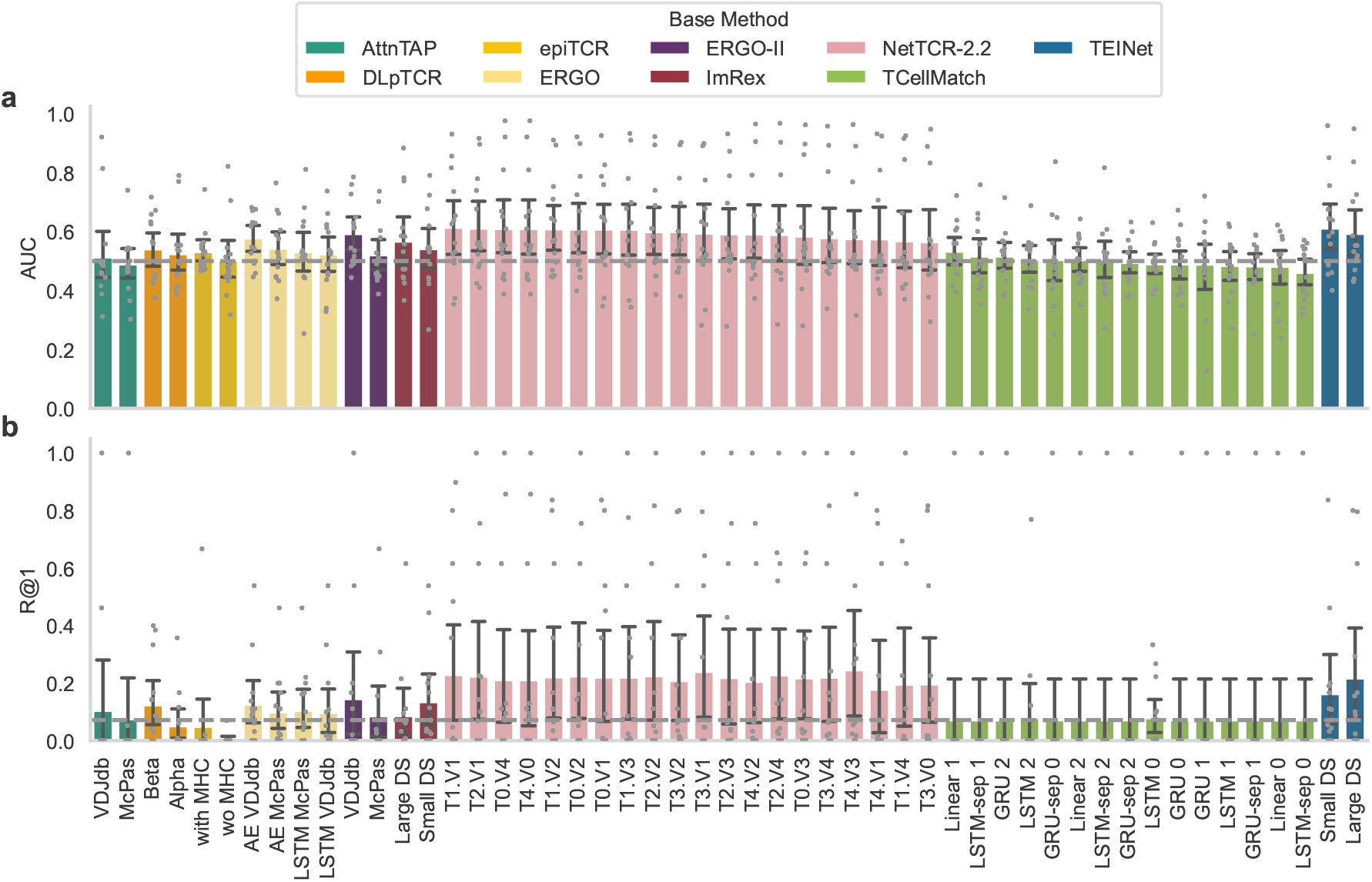
Performance of model alternatives on the viral dataset. Performance scores of methods with multiple model versions on the viral benchmark for AUC (**a**) and R@1 (**b**). The models are ordered by their performance on the respective metric within their method. The mean indicates the average metric score measured for each epitope individually, while the error bars indicate the 95% confidence interval over the scores (*n* = 14 epitopes). The dashed black line marks random predictions.

**Supplementary Figure 3.**
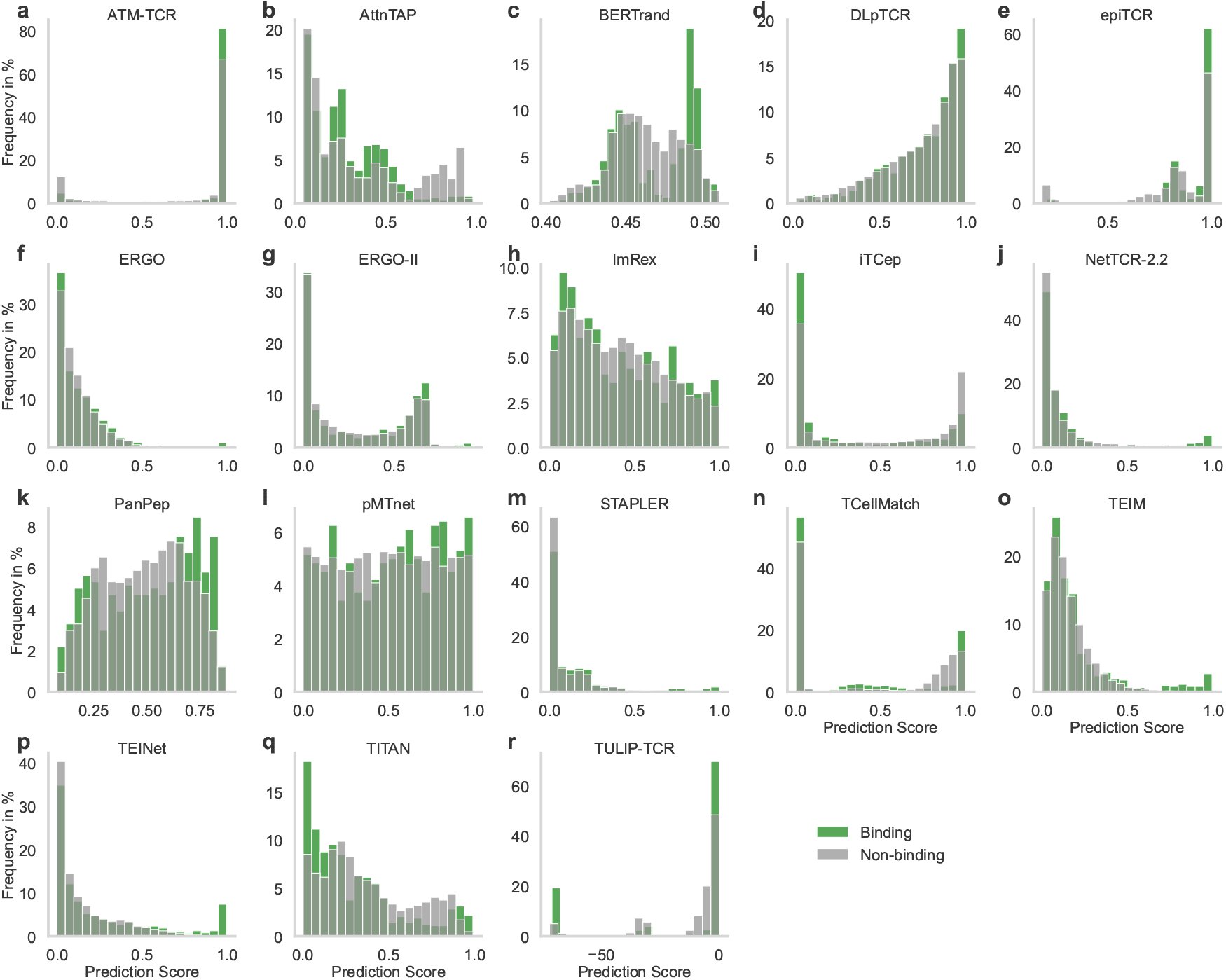
Distribution of prediction scores on the viral dataset. Binding score frequency for *n* = 20 bins between positive and negative pairs for all models (**a**-**r**).

**Supplementary Figure 4.**
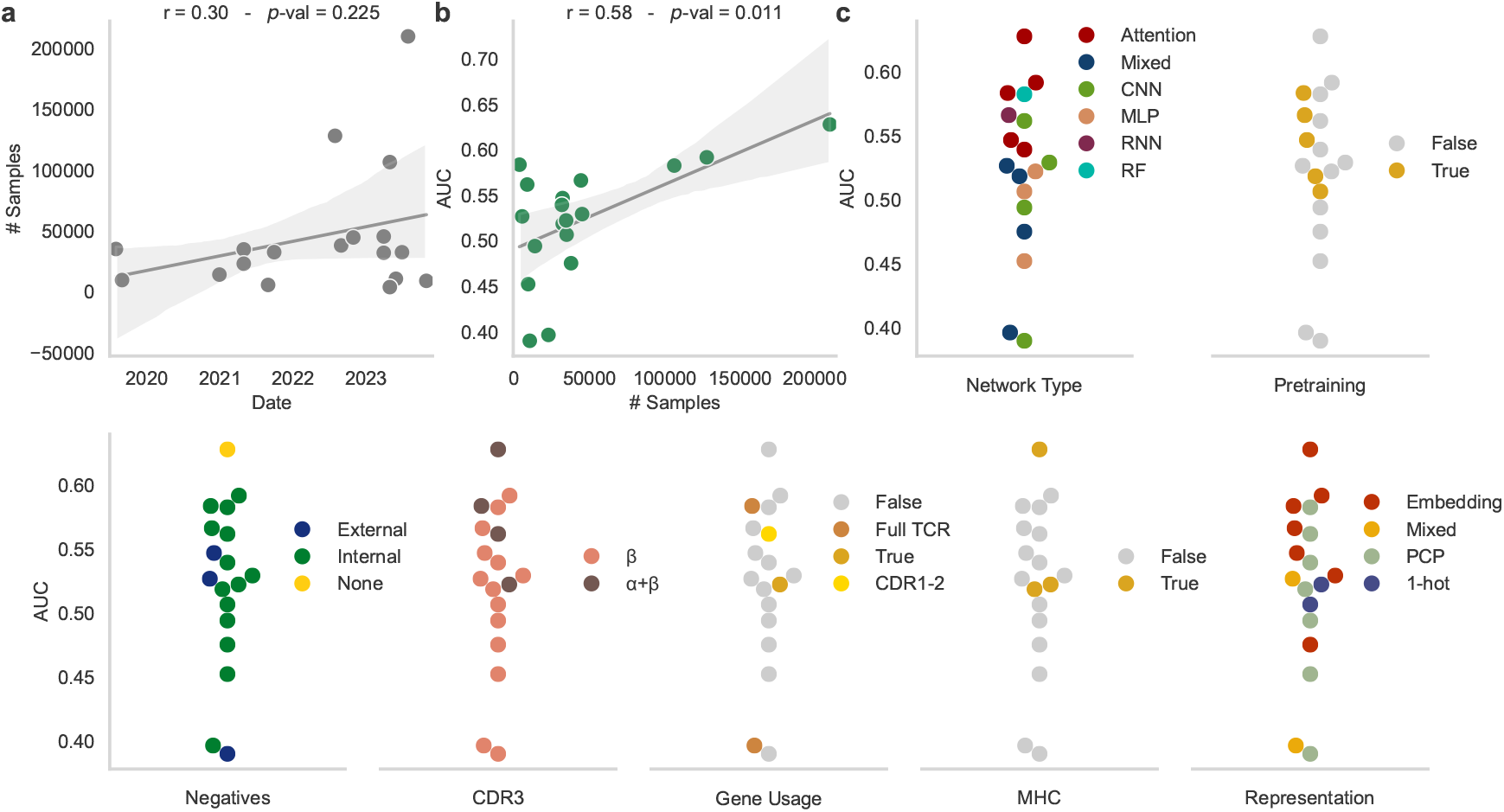
Impact of model properties on the performance on the viral dataset. **a**, Pearson correlation between the number of positive training samples and the initial publication date of the methods (*n* = 18). **b**, Pearson correlation between the average AUC score of the predictors and the number of positive training samples (*n* = 18). **c**, Average AUC indicated for different method design choices.

**Supplementary Figure 5.**
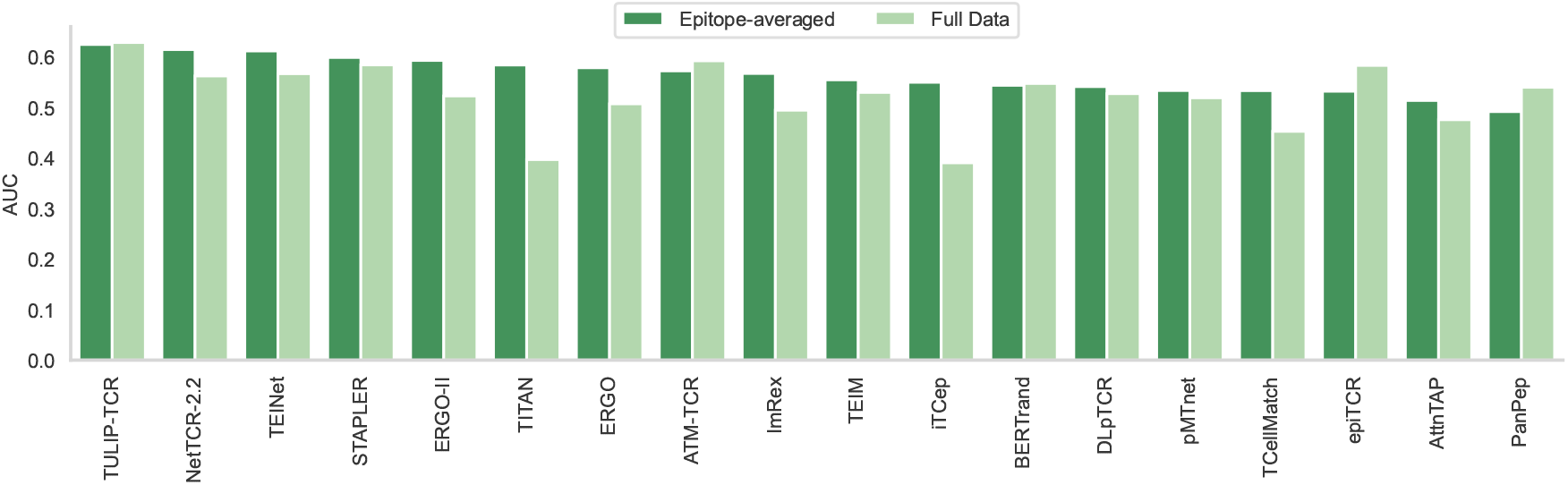
Differences in AUC calculation on the viral dataset. Comparison between the AUC averaged per epitope (*n* = 14) and the AUC calculated on the full dataset. The models are ordered by best averaged AUC.

**Supplementary Figure 6.**
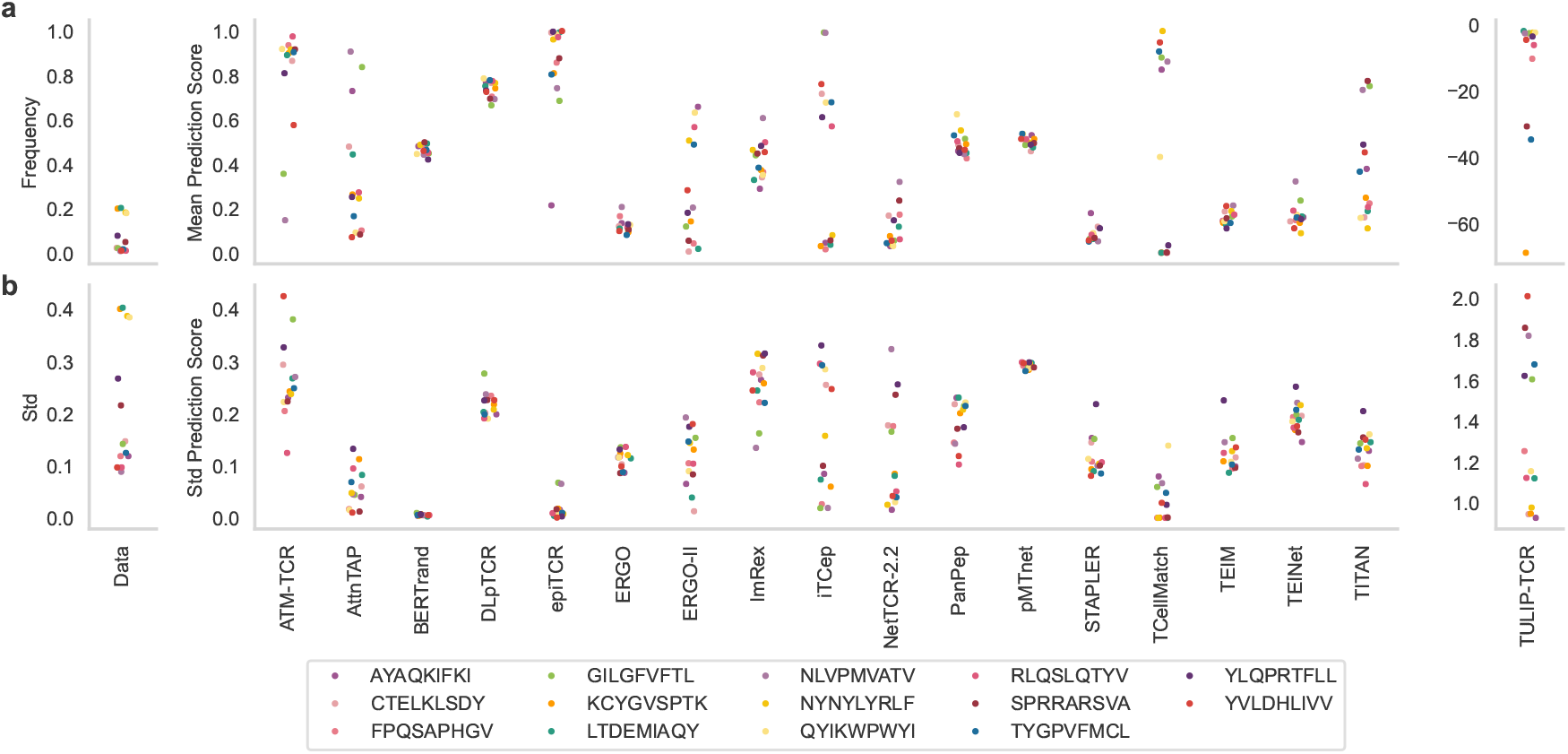
Prediction scores per epitope on the viral dataset. True data distribution and prediction scores statistics over all TCRs (*n* = 638) for each epitope indicating the mean (**a**) and standard deviation (**b**).

**Supplementary Figure 7.**
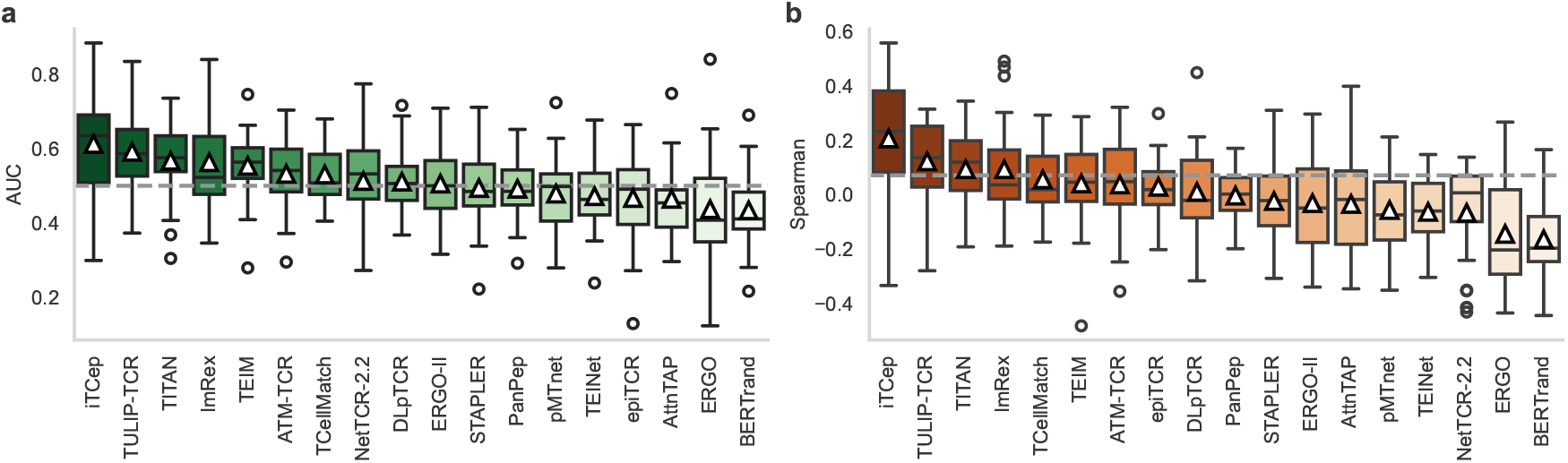
Performance of all models on the mutation dataset. Performance scores of all models on the mutation benchmark for AUC (**a**) and Spearman coefficient (**b**). The models are ordered by their performance on the respective metric. The box plot indicates the data quartiles while the median is indicated as a horizontal line and the mean as a white triangle (*n* = 14 TCRs). Outliers are marked separately. The dashed black line marks random predictions.

**Supplementary Figure 8.**
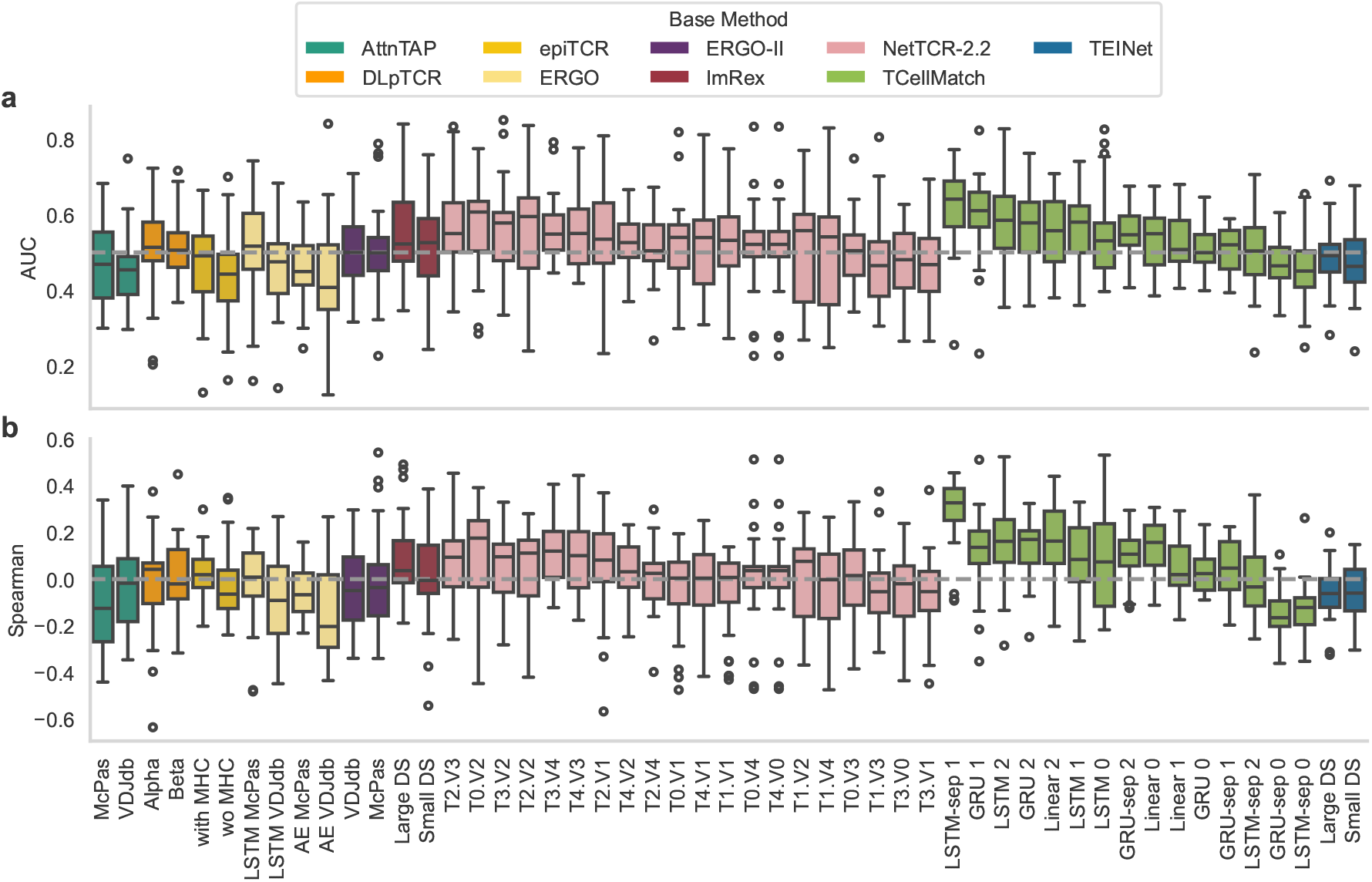
Performance of model alternatives on the mutation dataset. Performance scores of methods with multiple model versions on the mutation benchmark for AUC (**a**) and R@1 (**b**). The models are ordered by their performance on the respective metric within their method. The box plot indicates the data quartiles while the median is indicated as a horizontal line and the mean as a white triangle (*n* = 14 TCRs). Outliers are marked separately. The dashed black line marks random predictions.

**Supplementary Figure 9.**
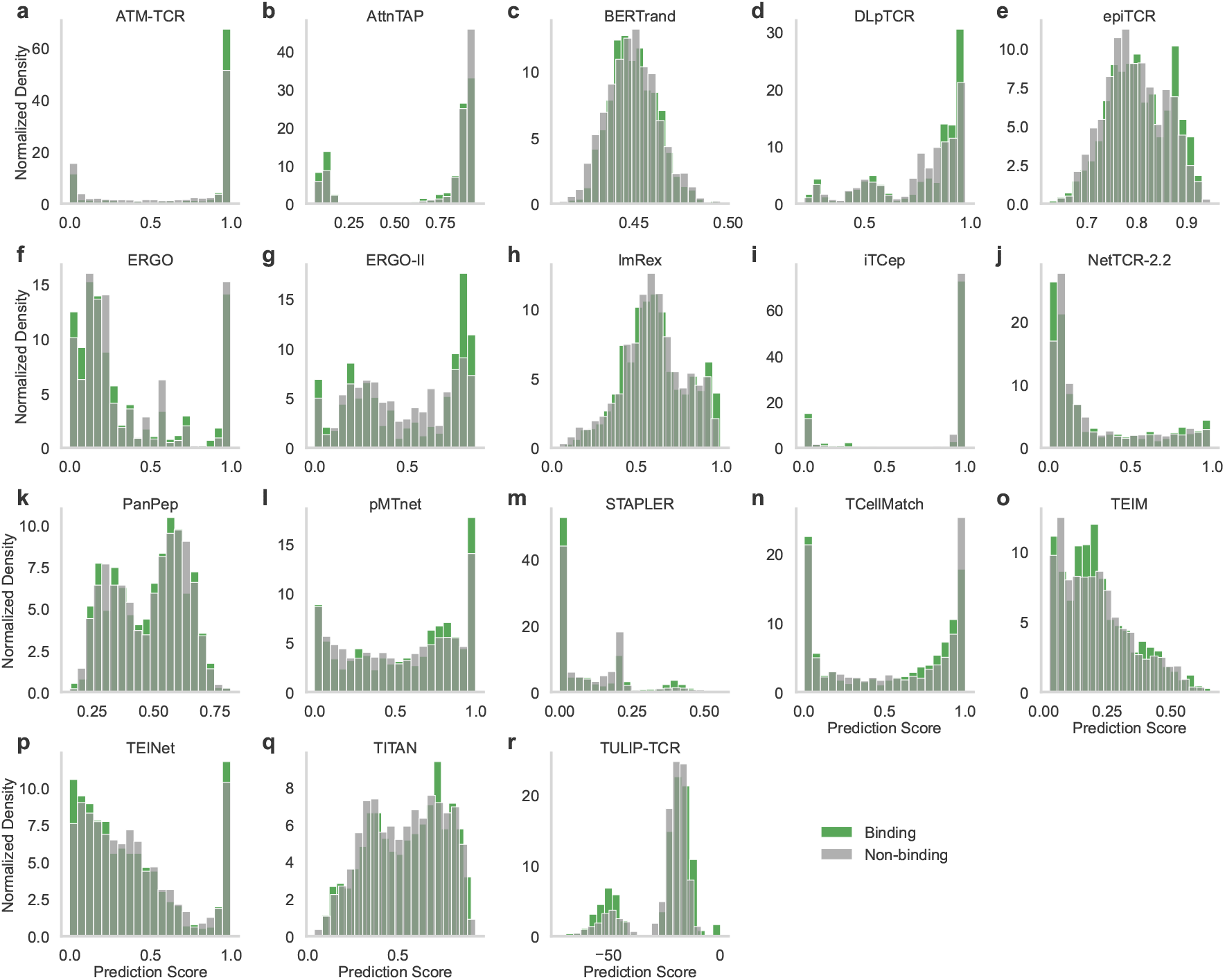
Distribution of prediction scores on the mutation dataset. Binding score frequency for *n* = 20 bins between positive and negative pairs for all models (**a**-**r**).

**Supplementary Figure 10.**
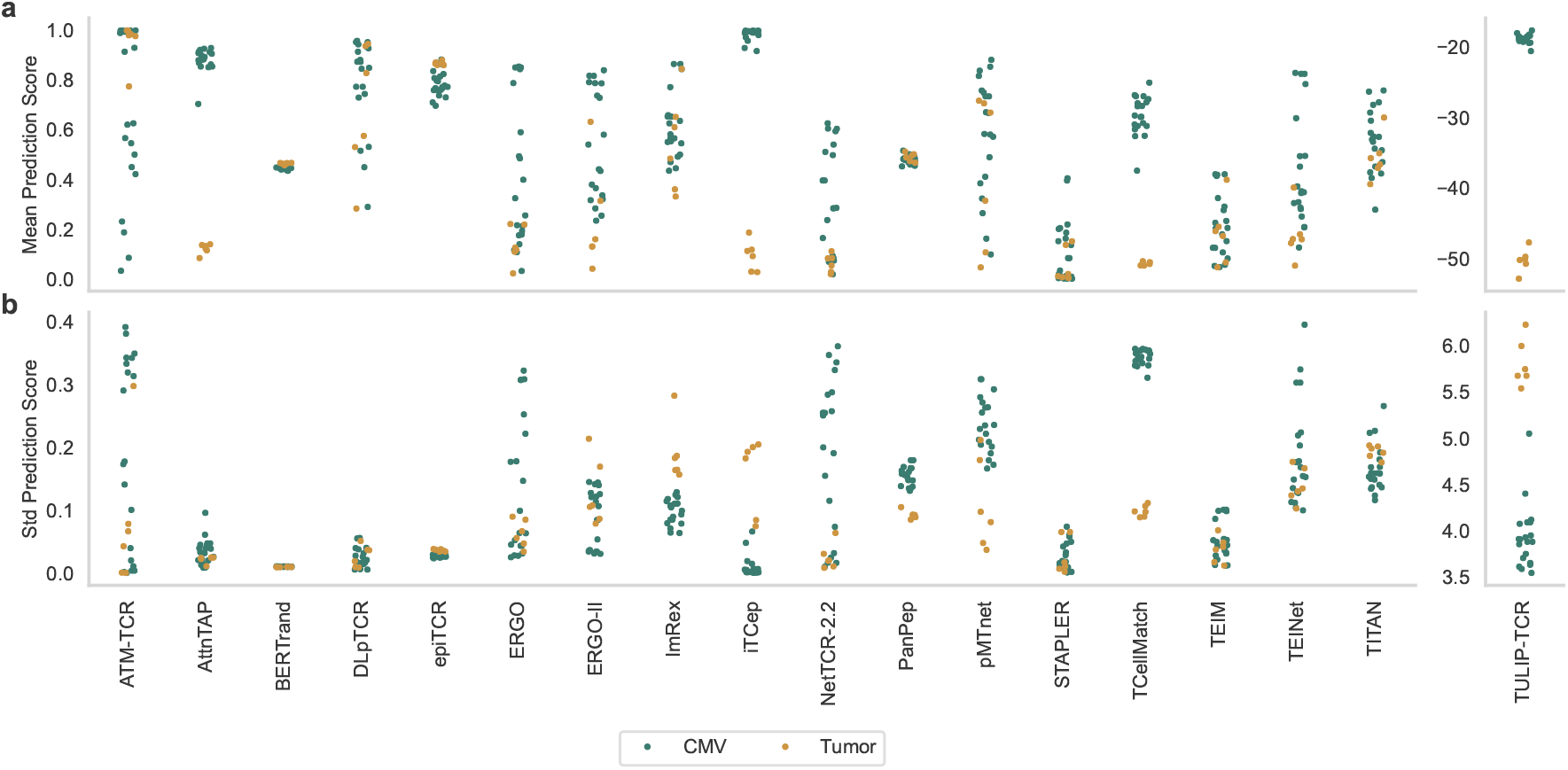
Prediction scores per epitope on the mutation dataset. True data distribution and prediction scores statistics over all epitopes (*n* = 134 for tumor TCRs, *n* = 172 for CMV TCRs) for each TCR indicating the mean (**a**) and standard deviation (**b**).

## Supplementary Tables

**Supplementary Table 1.** Additional predictor information. For each supported tool the name, the code source, first author, publication date, and license of the model is provided. The code repository of each predictor is linked via the name. The first occurrence in a journal, conference, or preprint is considered as the publication date.

| Name | Authors | Date | License | Reference |
| --- | --- | --- | --- | --- |
| ATM-TCR | Cai et al. | 07/2022 | Attribution 4.0 International | [35] |
| AttnTAP | Xu et al. | 08/2022 | Custom, non-commercial | [36] |
| BERtrand | Myronov et al. | 06/2023 | MIT | [37] |
| DLpTCR | Xu et al. | 08/2021 | Custom, non-commercial | [34] |
| epiTCR | Pham et al. | 04/2023 | MIT | [38] |
| ERGO | Springer et al. | 07/2019 | MIT | [17] |
| ERGO-II | Springer et al. | 04/2021 | MIT | [1] |
| ImRex | Moris et al. | 12/2020 | Custom, non-commercial | [18] |
| iTcep | Zhang et al. | 05/2023 | GNU | [39] |
| NetTCR-2.2 | Jensen et al. | 10/2023 | Custom, non-commercial | [40] |
| PanPep | Gao et al. | 03/2023 | GNU | [41] |
| pMTnet | Lu et al. | 09/2021 | GNU | [33] |
| STAPLER | Kwee et al. | 04/2023 | Apache License | [42] |
| TCellMatch | Fischer et al. | 08/2019 | BSD 3-Clause | [19] |
| TEIM | Peng et al. | 03/2023 | MIT | [43] |
| TEINet | Jiang et al. | 10/2022 | GNU | [44] |
| TITAN | Weber et al. | 04/2021 | Custom | [45] |
| TULIP-TCR | Meynard-Piganeau et al. | 07/2023 | GNU | [46] |

**Supplementary Table 2.** Performance on the viral dataset. The reported values correspond to the performance evaluated on either the full dataset or per epitope for classification (AUC: Area under the receiver operating characteristic curve, APS: average precision score, F1-Score) or ranking (Rank: Average rank of the correct epitope, R@K: Recall at k). The mean performance and standard deviation are indicated when the performance was evaluated per epitope. The three best scores for each metric are highlighted in bold.

|  | Full Data |  |  |  |  |  | Per Epitope |  |  |  |  |  |
| --- | --- | --- | --- | --- | --- | --- | --- | --- | --- | --- | --- | --- |
|  | AUC | APS | F1-Score | Rank | R@1 | R@3 | AUC | APS | F1-Score | Rank | R@1 | R@3 |
| ATM-TCR | **0.59** | 0.09 | 0.16 | **6.28** | 0.08 | 0.28 | 0.57±0.16 | 0.13±0.13 | **0.15±0.14** | 6.86±2.01 | 0.09±0.11 | 0.26±0.18 |
| AttnTAP | 0.48 | 0.07 | 0.14 | 7.80 | 0.02 | 0.04 | 0.51±0.16 | 0.12±0.16 | 0.10±0.13 | 7.29±4.20 | 0.10±0.29 | 0.21±0.43 |
| BERTrand | 0.55 | 0.09 | **0.21** | 6.92 | 0.07 | **0.43** | 0.54±0.12 | 0.12±0.13 | 0.06±0.13 | 7.35±4.05 | 0.08±0.26 | 0.21±0.42 |
| DLpTCR | 0.53 | 0.08 | 0.14 | **6.20** | 0.13 | 0.30 | 0.54±0.11 | 0.08±0.08 | 0.13±0.13 | 6.85±2.40 | 0.12±0.15 | 0.25±0.21 |
| epiTCR | 0.58 | 0.08 | 0.17 | 6.46 | 0.01 | 0.18 | 0.53±0.07 | 0.07±0.08 | 0.08±0.13 | 7.46±4.13 | 0.05±0.18 | 0.20±0.35 |
| ERGO | 0.51 | 0.09 | 0.13 | 7.45 | 0.09 | 0.25 | 0.58±0.09 | 0.13±0.14 | 0.13±0.12 | 6.36±1.71 | 0.13±0.15 | 0.32±0.15 |
| ERGO-II | 0.52 | 0.09 | 0.14 | 7.35 | 0.03 | 0.28 | 0.59±0.11 | 0.13±0.12 | 0.13±0.14 | 6.78±4.00 | 0.15±0.29 | 0.30±0.36 |
| ImRex | 0.49 | 0.08 | 0.13 | 7.68 | 0.09 | 0.24 | 0.57±0.15 | 0.12±0.12 | 0.12±0.13 | 7.12±2.09 | 0.09±0.16 | 0.28±0.26 |
| iTCep | 0.39 | 0.05 | 0.13 | 8.93 | 0.02 | 0.10 | 0.55±0.12 | 0.11±0.14 | 0.12±0.13 | 7.23±3.93 | 0.09±0.26 | 0.25±0.37 |
| NetTCR-2.2 | 0.56 | 0.11 | 0.15 | 7.02 | 0.13 | **0.23** | **0.61±0.17** | 0.17±0.22 | 0.14±0.13 | **5.97±3.05** | **0.23±0.33** | 0.34±0.38 |
| PanPep | 0.54 | 0.09 | 0.16 | 6.59 | 0.13 | **0.32** | 0.49±0.10 | 0.09±0.12 | 0.09±0.12 | 7.60±2.60 | 0.07±0.19 | 0.20±0.27 |
| pMTnet | 0.52 | 0.08 | 0.14 | 7.27 | 0.08 | 0.24 | 0.53±0.06 | 0.10±0.10 | 0.12±0.12 | 6.93±1.06 | 0.10±0.14 | 0.28±0.12 |
| STAPLER | **0.58** | **0.15** | 0.16 | 6.42 | 0.13 | 0.31 | 0.60±0.17 | **0.19±0.25** | 0.15±0.14 | **5.90±2.19** | **0.22±0.30** | **0.37±0.32** |
| TCellMatch | 0.45 | 0.08 | 0.18 | 8.11 | **0.18** | 0.21 | 0.53±0.09 | 0.08±0.08 | 0.02±0.08 | 7.50±4.18 | 0.07±0.27 | 0.21±0.43 |
| TEIM | 0.53 | 0.14 | 0.15 | 7.03 | 0.12 | 0.26 | 0.55±0.18 | 0.17±0.24 | 0.12±0.22 | 6.25±3.00 | 0.18±0.28 | **0.38±0.36** |
| TEINet | 0.57 | **0.19** | **0.19** | 6.81 | **0.16** | 0.32 | **0.61±0.16** | **0.20±0.24** | **0.16±0.22** | **6.04±2.46** | 0.16±0.23 | 0.36±0.24 |
| TITAN | 0.40 | 0.07 | 0.13 | 8.88 | 0.09 | 0.16 | 0.58±0.15 | 0.15±0.17 | 0.12±0.13 | 6.71±3.78 | 0.16±0.28 | 0.29±0.44 |
| TULIP-TCR | **0.63** | **0.25** | **0.26** | **6.00** | **0.26** | **0.44** | **0.62±0.17** | **0.22±0.26** | **0.15±0.24** | 6.42±4.57 | **0.24±0.30** | **0.38±0.38** |

**Supplementary Table 3.** Performance on the mutation dataset. The reported values correspond to the performance evaluated on either the full dataset or per TCR for classification (AUC: Area under the receiver operating characteristic curve, APS: average precision score, F1-Score) or regression (Pearson and Spearman correlation coefficient). The mean performance and standard deviation are indicated when the performance was evaluated per TCR. The three best scores for each metric are highlighted in bold.

|  | Full Data |  |  |  |  | Per TCR |  |  |  |  |
| --- | --- | --- | --- | --- | --- | --- | --- | --- | --- | --- |
|  | AUC | APS | F1-Score | Pearson | Spearman | AUC | APS | F1-Score | Pearson | Spearman |
| ATM-TCR | 0.50 | 0.31 | **0.48** | **0.07** | -0.05 | 0.53±0.10 | **0.34±0.17** | 0.37±0.22 | 0.06±0.16 | 0.04±0.16 |
| AttnTAP | 0.47 | 0.29 | 0.47 | -0.04 | -0.03 | 0.46±0.11 | 0.29±0.17 | 0.42±0.19 | -0.05±0.18 | -0.03±0.19 |
| BERTrand | 0.45 | 0.27 | 0.47 | -0.08 | -0.09 | 0.43±0.10 | 0.26±0.16 | 0.42±0.19 | -0.12±0.16 | -0.16±0.15 |
| DLpTCR | 0.54 | 0.35 | 0.47 | 0.06 | **0.10** | 0.51±0.09 | 0.29±0.16 | 0.31±0.27 | 0.03±0.15 | 0.01±0.16 |
| epiTCR | 0.53 | 0.32 | 0.47 | 0.04 | **0.10** | 0.47±0.11 | 0.29±0.17 | 0.41±0.19 | -0.02±0.13 | 0.03±0.11 |
| ERGO | 0.53 | 0.32 | **0.48** | 0.03 | 0.02 | 0.44±0.15 | 0.29±0.14 | 0.40±0.21 | -0.17±0.21 | -0.14±0.19 |
| ERGO-II | **0.54** | **0.37** | **0.48** | 0.03 | -0.01 | 0.51±0.09 | 0.32±0.19 | 0.28±0.29 | -0.01±0.20 | -0.03±0.17 |
| ImRex | 0.53 | **0.36** | 0.48 | 0.07 | 0.05 | 0.56±0.14 | 0.34±0.20 | **0.42±0.19** | **0.09±0.18** | 0.09±0.18 |
| iTCep | **0.56** | **0.38** | 0.47 | **0.09** | **0.11** | **0.61±0.15** | **0.39±0.19** | **0.42±0.19** | **0.09±0.19** | **0.21±0.22** |
| NetTCR-2.2 | 0.43 | 0.28 | 0.47 | -0.13 | -0.16 | 0.51±0.13 | 0.31±0.14 | 0.42±0.19 | -0.04±0.19 | -0.07±0.18 |
| PanPep | 0.48 | 0.30 | 0.47 | -0.03 | -0.04 | 0.49±0.08 | 0.31±0.18 | 0.41±0.19 | -0.01±0.09 | -0.00±0.08 |
| pMTnet | **0.54** | 0.34 | 0.47 | **0.09** | 0.09 | 0.48±0.10 | 0.30±0.16 | 0.41±0.19 | -0.03±0.13 | -0.06±0.16 |
| STAPLER | 0.47 | 0.32 | 0.47 | -0.02 | -0.10 | 0.49±0.10 | 0.30±0.18 | 0.42±0.19 | -0.00±0.11 | -0.02±0.17 |
| TCellMatch | 0.54 | 0.32 | 0.47 | 0.07 | 0.09 | 0.53±0.07 | 0.29±0.16 | 0.42±0.17 | 0.07±0.08 | 0.06±0.14 |
| TEIM | 0.48 | 0.29 | 0.47 | -0.03 | -0.02 | 0.55±0.09 | 0.33±0.18 | 0.38±0.22 | 0.05±0.17 | 0.04±0.16 |
| TEINet | 0.52 | 0.33 | 0.47 | 0.04 | 0.03 | 0.47±0.09 | 0.29±0.14 | 0.41±0.19 | -0.07±0.15 | -0.06±0.13 |
| TITAN | 0.52 | 0.32 | 0.47 | 0.01 | -0.00 | **0.57±0.10** | 0.33±0.16 | 0.42±0.19 | 0.08±0.15 | **0.09±0.14** |
| TULIP-TCR | 0.53 | 0.34 | 0.48 | 0.05 | 0.01 | **0.59±0.11** | **0.40±0.14** | **0.42±0.18** | **0.16±0.18** | **0.12±0.17** |

---

**Supplementary Data 1** | **Benchmarking results on the viral dataset**. All metric scores (AUC, APS, F1-Score, average Rank, R@1, R@3) on the full data, macro-averaged, averaged, and per epitope.

**Supplementary Data 2** | **Predictions the viral dataset**. Prediction score for each method between all combinations of epitopes and TCRs of the viral dataset.

**Supplementary Data 3** | **Benchmarking results on the mutation dataset**. All metric scores (AUC, APS, F1-Score, Pearson correlation, Spearman coefficient) on the full data, macro-averaged, averaged, and per TCR.

**Supplementary Data 4** | **Predictions the mutation dataset**. Prediction score for each method between all combinations of epitopes and TCRs of the mutation dataset.

## References

[1] Springer, I., Tickotsky, N. & Louzoun, Y. Contribution of t cell receptor alpha and beta cdr3, mhc typing, v and j genes to peptide binding prediction. Frontiers in immunology 12, 664514 (2021).

[2] Davis, M. M. T cell analysis in vaccination. Current opinion in immunology 65, 70–73 (2020).

[3] D’Ippolito, E., Wagner, K. I. & Busch, D. H. Needle in a haystack: the naïve repertoire as a source of t cell receptors for adoptive therapy with engineered t cells. International Journal of Molecular Sciences 21, 8324 (2020).

[4] Serr, I., Drost, F., Schubert, B. & Daniel, C. Antigen-specific treg therapy in type 1 diabetes– challenges and opportunities. Frontiers in immunology 12, 712870 (2021).

[5] Scott, D. & Singer, D. S. Harnessing the power of discovery. Cancer Discovery 13, 819–823 (2023).

[6] 10x Genomics. A new way of exploring immunity–linking highly multiplexed antigen recognition to immune repertoire and phenotype. Tech. rep (2019).

[7] Dash, P. et al. Quantifiable predictive features define epitope-specific t cell receptor repertoires. Nature 547, 89–93 (2017).

[8] Chronister, W. D. et al. Tcrmatch: predicting t-cell receptor specificity based on sequence similarity to previously characterized receptors. Frontiers in immunology 12, 640725 (2021).

[9] Sidhom, J.-W., Larman, H. B., Pardoll, D. M. & Baras, A. S. Deeptcr is a deep learning framework for revealing sequence concepts within t-cell repertoires. Nature communications 12, 1605 (2021).

[10] Wu, K. et al. Tcr-bert: learning the grammar of t-cell receptors for flexible antigen-xbinding analyses. Biorxiv 2021–11 (2021).

[11] Drost, F., Schiefelbein, L. & Schubert, B. metcrs-learning a metric for t-cell receptors. BioRxiv 2022–10 (2022).

[12] Vita, R. et al. The immune epitope database (iedb): 2018 update. Nucleic acids research 47, D339–D343 (2019).

[13] Bagaev, D. V. et al. Vdjdb in 2019: database extension, new analysis infrastructure and a t-cell receptor motif compendium. Nucleic Acids Research 48, D1057–D1062 (2020).

[14] Tickotsky, N., Sagiv, T., Prilusky, J., Shifrut, E. & Friedman, N. Mcpas-tcr: a manually curated catalogue of pathology-associated t cell receptor sequences. Bioinformatics 33, 2924– 2929 (2017).

[15] Jokinen, E., Huuhtanen, J., Mustjoki, S., Heinonen, M. & Lähdesmäki, H. Predicting recognition between t cell receptors and epitopes with tcrgp. PLoS computational biology 17, e1008814 (2021).

[16] Jurtz, V. I. et al. Nettcr: sequence-based prediction of tcr binding to peptide-mhc complexes using convolutional neural networks. BioRxiv 433706 (2018).

[17] Springer, I., Besser, H., Tickotsky-Moskovitz, N., Dvorkin, S. & Louzoun, Y. Prediction of specific tcr-peptide binding from large dictionaries of tcr-peptide pairs. Frontiers in immunology 11, 1803 (2020).

[18] Moris, P. et al. Current challenges for unseen-epitope tcr interaction prediction and a new perspective derived from image classification. Briefings in Bioinformatics 22, bbaa318 (2021).

[19] Fischer, D. S., Wu, Y., Schubert, B. & Theis, F. J. Predicting antigen specificity of single t cells based on tcr cdr 3 regions. Molecular systems biology 16, e9416 (2020).

[20] Deng, L. et al. Performance comparison of tcr-pmhc prediction tools reveals a strong data dependency. Frontiers in Immunology 14, 1128326 (2023).

[21] Dens, C., Laukens, K., Bittremieux, W. & Meysman, P. The pitfalls of negative data bias for the t-cell epitope specificity challenge. bioRxiv 2023–04 (2023).

[22] Meysman, P. et al. Benchmarking solutions to the t-cell receptor epitope prediction problem: Immrep22 workshop report. ImmunoInformatics 9, 100024 (2023).

[23] Barton, J. Immrep23: Tcr specificity prediction challenge (2023).

[24] Nielsen, M. et al. Lessons learned from the immrep23 tcr-epitope prediction challenge. ImmunoInformatics (2024).

[25] Schubert, B. et al. Fred 2: an immunoinformatics framework for python. Bioinformatics 32, 2044–2046 (2016).

[26] Kocher, K. et al. Quality of vaccination-induced t cell responses is conveyed by polyclonality and high, but not maximum, antigen receptor avidity. bioRxiv 2024–10 (2024).

[27] Habern, O. Introducing beam (barcode enabled antigen mapping): Benefits of rapid, antigenspecific b-and t-cell discovery. 10x Genomics (2022).

[28] Drost, F. et al. Predicting t cell receptor functionality against mutant epitopes. Cell Genomics 4 (2024).

[29] Vander Heiden, J. A. et al. Airr community standardized representations for annotated immune repertoires. Frontiers in immunology 9, 2206 (2018).

[30] Sturm, G. et al. Scirpy: a scanpy extension for analyzing single-cell t-cell receptor-sequencing data. Bioinformatics 36, 4817–4818 (2020).

[31] Zhang, W. et al. Pird: pan immune repertoire database. Bioinformatics 36, 897–903 (2020).

[32] Francis, J. M. et al. Allelic variation in class i hla determines cd8+ t cell repertoire shape and cross-reactive memory responses to sars-cov-2. Science immunology 7, eabk3070 (2021).

[33] Lu, T. et al. Deep learning-based prediction of the t cell receptor–antigen binding specificity. Nature machine intelligence 3, 864–875 (2021).

[34] Xu, Z. et al. Dlptcr: an ensemble deep learning framework for predicting immunogenic peptide recognized by t cell receptor. Briefings in Bioinformatics 22, bbab335 (2021).

[35] Cai, M., Bang, S., Zhang, P. & Lee, H. Atm-tcr: Tcr-epitope binding affinity prediction using a multi-head self-attention model. Frontiers in Immunology 13, 893247 (2022).

[36] Xu, Y. et al. Attntap: A dual-input framework incorporating the attention mechanism for accurately predicting tcr-peptide binding. Frontiers in Genetics 13, 942491 (2022).

[37] Myronov, A., Mazzocco, G., Krol, P. & Plewczynski, D. Bertrand-peptide: Tcr binding prediction using bidirectional encoder representations from transformers augmented with random tcr pairing. bioRxiv 2023–06 (2023).

[38] Pham, M.-D. N. et al. epitcr: a highly sensitive predictor for tcr–peptide binding. Bioinformatics 39, btad284 (2023).

[39] Zhang, Y. et al. itcep: a deep learning framework for identification of t cell epitopes by harnessing fusion features. Frontiers in Genetics 14, 1141535 (2023).

[40] Jensen, M. F. & Nielsen, M. Nettcr 2.2-improved tcr specificity predictions by combining pan-and peptide-specific training strategies, loss-scaling and integration of sequence similarity. bioRxiv 2023–10 (2023).

[41] Gao, Y. et al. Pan-peptide meta learning for t-cell receptor–antigen binding recognition. Nature Machine Intelligence 5, 236–249 (2023).

[42] Kwee, B. P. et al. Stapler: Efficient learning of tcr-peptide specificity prediction from full-length tcr-peptide data. bioRxiv 2023–04 (2023).

[43] Peng, X. et al. Characterizing the interaction conformation between t-cell receptors and epitopes with deep learning. Nature Machine Intelligence 5, 395–407 (2023).

[44] Jiang, Y., Huo, M. & Cheng Li, S. Teinet: a deep learning framework for prediction of tcr– epitope binding specificity. Briefings in Bioinformatics 24, bbad086 (2023).

[45] Weber, A., Born, J. & Rodriguez Martínez, M. Titan: T-cell receptor specificity prediction with bimodal attention networks. Bioinformatics 37, i237–i244 (2021).

[46] Meynard-Piganeau, B., Feinauer, C., Weigt, M., Walczak, A. M. & Mora, T. Tulip-a transformer based unsupervised language model for interacting peptides and t-cell receptors that generalizes to unseen epitopes. bioRxiv 2023–07 (2023).

[47] Weininger, D., Weininger, A. & Weininger, J. L. Smiles. 2. algorithm for generation of unique smiles notation. Journal of chemical information and computer sciences 29, 97–101 (1989).

[48] Grazioli, F. et al. On tcr binding predictors failing to generalize to unseen peptides. Frontiers in immunology 13, 1014256 (2022).

[49] Heather, J. M. et al. Stitchr: stitching coding tcr nucleotide sequences from v/j/cdr3 information. Nucleic Acids Research 50, e68–e68 (2022).

[50] Dunbar, J. & Deane, C. M. Anarci: antigen receptor numbering and receptor classification. Bioinformatics 32, 298–300 (2016).

[51] Kocher, K. et al. Adaptive immune responses are larger and functionally preserved in a hypervaccinated individual. The Lancet Infectious Diseases 24, e272–e274 (2024).

[52] Straub, A. et al. Recruitment of epitope-specific t cell clones with a low-affinity threshold supports efficacy against mutational escape upon re-infection. Immunity (2023).

[53] Oh Song, H., Xiang, Y., Jegelka, S. & Savarese, S. Deep metric learning via lifted structured feature embedding. In Proceedings of the IEEE conference on computer vision and pattern recognition, 4004–4012 (2016).

